# Dense Longitudinal Precision Neuroimaging of Recovery from Traumatic Brain Injury

**DOI:** 10.1101/2025.03.09.642218

**Authors:** Aishwarya Rajesh, Timothy O. Laumann, Deanna Barch, Benjamin P. Kay, Ashley C. Meyer, Terrance T. Kummer, Nico U.F. Dosenbach, Evan M. Gordon

## Abstract

Traumatic brain injury (TBI) disrupts white matter tracts essential for cognition and emotion. Diffusion tensor imaging (DTI) can noninvasively measure white matter integrity. However, DTI has been inconsistent in predicting patient recovery from TBI, possibly due to the complex, dynamic, and individual-specific process of post-TBI white matter remodeling. Here, we employed dense longitudinal neuroimaging to track white matter recovery weekly over six months after a TBI within a single patient and a control in a similar age group (21 vs. 24 y.o.). In the patient, but not in the control, DTI metrics precisely tracked parabolic trajectories across time, with early structural alterations continuing for more than 15 weeks before reversing direction. The extent of alteration in each tract was correlated with the time until reversal. These continuous DTI changes also mediated recovery of cognitive and emotional function, suggesting they are not passive markers of damage but dynamic processes underlying functional improvement. Complementary diffusion basis spectrum imaging (DBSI) revealed an initial phase of cellular loss followed by inflammatory remodeling, vascular adaptations, and persistent metabolic activity. Our findings indicate that recovery does not follow predefined phases but rather individualized transition points, which could define optimal windows for rehabilitation. Identifying these inflection points may enable personalized interventions aligned with biologically relevant structural shifts, rather than broad recovery periods.

## Introduction

Nearly half of the global population is expected to experience a traumatic brain injury (TBI) in their lifetime (Jiang et al., 2023; Maas et al., 2022). In TBI, shearing forces during head impact disrupt the microstructural integrity of white matter tracts (Gennarelli et al., 1982). These tracts connect distant brain regions that support cognition and behavior (Schmahmann et al., 2008, Filley 2012, Catani et al., 2012, Forkel et al., 2021, Roberts, Anderson, & Husain, 2013). Thus, understanding how white matter integrity relates to functional impairment is critical for diagnosis and prognosis of TBI patients.

White matter damage induced by TBI is best assessed using diffusion tensor imaging (DTI), a noninvasive neuroimaging technique that measures water diffusion in the brain (Hulkower et al., 2013). In healthy white matter, diffusion is anisotropic, occurring primarily along axonal pathways due to the restrictive influence of myelin sheaths and cellular structures. Disruption of these tracts results in more isotropic diffusion. Fractional Anisotropy (FA)—the most common DTI metric—indexes the degree of isotropic diffusion and is consistently reduced (implying more white matter disruption) in individuals with TBI, compared to healthy controls (Nakayama et al, 2006, Hulkower et al., 2013, Niogi & Pratik, 2010).

TBI is a dynamic rather than a static condition, as it involves an extended post-injury recovery period, during which impaired cognitive and emotional processes generally improve (Caroll et al., 2004; Shretlen & Shapiro, 2003; Frencham, Fox, & Maybery, 2005). Here, we define the post-TBI recovery period as the interval following acute stabilization, extending through subacute and chronic phases of recovery, during which white matter remodeling and functional improvements continue to evolve.

However, recovery trajectories vary widely across individuals, and there is little consensus on precise time boundaries, with studies defining subacute recovery as anywhere from one week to three months post-injury and chronic recovery extending beyond three months (Amyot et al., 2015; Wallace, Mathias, & Ward, 2018). Ideally, DTI metrics would provide mechanistic and clinically predictive power for understanding the timing and extent of this ongoing recovery process. Unfortunately, while DTI metrics differ between controls and TBI patients, their relationship with TBI recovery is less clear. Prior work has argued that FA measures increase (Chiou et al., 2019; Ling et al., 2012; Mayer, 2010), decrease (Niogi & Pratik, 2010; Veeramuthu et al., 2016; Næss-Schmidt et al., 2017; Karlsen et al., 2019; Palacios et al., 2020), or remain unchanged (Eirud et al., 2014; Narayana et al., 2015; Wilde et al., 2016) during this period (Kim et al., 2022). Similarly, while greater alterations in DTI metrics are cross-sectionally associated with cognitive deficits (Preziosa et al., 2022; Gyebnár et al., 2018; Mesaros et al., 2012; Brickman et al., 2006) and increased susceptibility to mood disturbances (Sexton, Mackay, & Ebmeier, 2009; Murphy & Frodl, 2011; Catani et al., 2012; Zhang et al., 2011; Han et al., 2008), it remains unclear whether recovery from these deficits relates to ongoing changes in DTI metrics.

Study design limitations have prevented DTI from becoming a reliable tool for tracking TBI recovery in clinical settings. First, most DTI studies tracking post-TBI recovery collect only two to three post-injury timepoints (Kim et al., 2022). This sparse sampling limits the ability to track the dynamics of white matter changes over the course of recovery or to establish associations with the resolution of cognitive and emotional symptoms. Second, most DTI studies rely on group-level designs that average across patients and brain regions, but this approach overlooks the substantial heterogeneity in injury characteristics and recovery trajectories (Rosenbaum & Lipton, 2012, Kim et al., 2022; Eirud et al., 2014). As a result, such studies may lack the precision needed to capture meaningful recovery-related changes. In contrast, examining detailed longitudinal data from an individual can provide a clearer view of within-person recovery dynamics, potentially revealing patterns obscured by group averaging. Furthermore, although single-case studies do not offer broad generalizability, they can generate hypotheses and mechanistic insights that larger studies can later test. Third, recent work has argued that effects observed at the group level—and particularly human neuroimaging measures—may be fundamentally unsuited to use for inferring within-individual processes (Mattoni et al., 2025). For instance, group-level findings (e.g., lower FA is associated with worse outcomes) may not translate to within-person processes (e.g., an individual’s FA might first decrease but later recover alongside cognitive improvements). Thus, while reliable group differences in DTI measures between TBI patients and controls can offer valuable insights, they may not directly translate to mechanisms of within-individual TBI recovery. To overcome this limitation, approaches that capture recovery dynamics at the individual level—such as “precision imaging”— are necessary (Gordon et al., 2017).

Precision imaging improves the reliability of neuroimaging measures by repeatedly scanning individuals over time, reducing measurement error, and eliminating confounds introduced by cross-individual variability in brain function and organization (Gordon et al., 2017; Laumann et al., 2015; Gordon et al., 2023; Gratton, Nelson, & Gordon, 2022; Braga & Buckner, 2017; Naselaris et al., 2021, Michon et al., 2022).

Precision imaging approaches are argued to be critical for clinical translation of neuroimaging findings (Gratton et al., 2020; Kraus et al., 2023; Laumann et al., 2023; Mattoni et al., 2025). Precision imaging has been particularly well suited for examining longitudinal within-individual associations between neuroimaging measures and variable external processes, including mood symptoms (Lynch et al., 2024), motor behavior (Newbold et al., 2020), effects of psychedelic drugs (Siegel et al., 2024), hormonal influences of the menstrual cycle (Pritschet et al., 2020) and pregnancy (Pritschet et al., 2024). While many precision imaging studies include multiple individuals to allow for some level of generalizability (e.g., Gordon et al., 2023, Lynch et al., 2024), the strength of this approach lies in its ability to capture meaningful within-person associations across time —an approach that has been particularly well demonstrated in single-subject studies (e.g., Pritschet et al., 2024). This emphasis on within-person associations is especially crucial for studying longitudinal recovery-related changes in DTI measures following TBI, where group-level approaches may obscure individual recovery trajectories. Thus, to capture these individualized recovery dynamics, dense longitudinal precision imaging is necessary for understanding how post-TBI changes in DTI measures track cognitive and emotional recovery, with single-subject approaches offering a critical lens that group-level analyses cannot resolve.

To this end, we used an intensive, densely sampled, longitudinal precision imaging approach to systematically track individual-level white matter changes following TBI in a single TBI patient, as well as in a healthy, non-brain injured control in a similar age group (21 vs. 24 y.o.). We scanned the patient weekly beginning two weeks post-injury and continuing for six months, with a 1-year follow-up, to characterize the post-TBI trajectory of DTI metrics and to examine the relationship between DTI metrics and cognitive and emotional outcomes. Importantly, DTI metrics like FA reflect not only the degree of intact anisotropic diffusion, but also dynamic secondary injury processes such as inflammation, edema, and axonal repair (Armstrong et al., 2016, Palacios et al., 2020); it is unclear which factors are most related to post-TBI symptomatic improvement. Thus, we further employed Diffusion Basis Spectrum Imaging (Wang et al., 2019a, 2015, 2011), an advanced technique that can disentangle these overlapping processes, to improve specificity in characterizing TBI-related white matter changes.

The dense repeated imaging collected for this study enables an unprecedented, integrated analysis of white matter recovery trajectories and their association with behavioral outcomes. It thus represents a critical step forward in defining how white matter integrity evolves post-TBI and in laying the groundwork for biomarker-driven, individualized rehabilitation strategies.

## Results

### 1. Nonlinear White Matter Trajectories Reveal Dynamic Post-TBI Remodeling

In both a participant with TBI and a control participant, 18 major white matter tracts were defined in each participant using TRACULA (Yendiki et al., 2011). DTI metrics—including fractional anisotropy (FA), radial diffusivity (RD), axial diffusivity (AD), and mean diffusivity (MD)—were extracted from each of these tracts over 27 consecutive weeks post-injury (or post enrollment, for the control participant).

We hypothesized that we would observe linear trajectories of DTI metrics over the course of the post-injury period in the TBI patient but not in the control. Remarkably, initial examination of these trajectories in the TBI patient suggested not a linear effect, but very tight adherence to a curvilinear trajectory, with an initial steep change, particularly in the FA, RD, and MD tracts (a decline for FA [Fig. 1]; an increase for RD and MD [Fig. S1]) followed by a gradual reversal in direction. This pattern was observed across many tracts (see Fig 1B, S1B, F for examples; Fig S2 for FA changes across all tracts).

**Figure 1:**
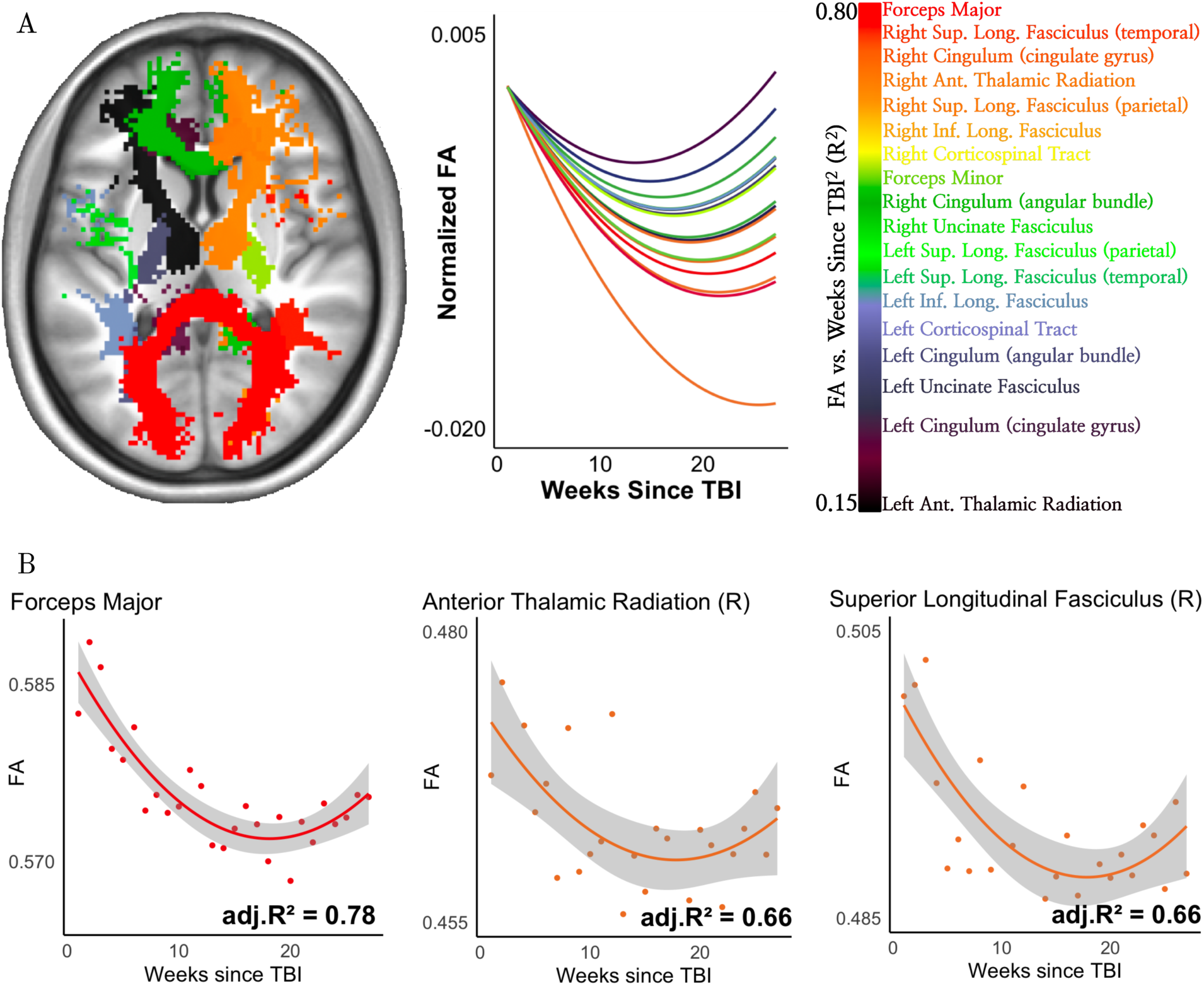
Nonlinear Trajectories of Fractional Anisotropy Changes (FA) Over Time Post- TBI. (A) Left: White matter tracts for TBI patient are colored by strength of model (quadratic) fit for FA over time (in weeks) since TBI. Tracts in orange-red indicate a strong relationship (variance explained ∼80%), while tracts in purple-black indicate a weaker relationship (variance explained ∼15%). Right: FA trajectories across all 18 tracts over time, based on the best-fitting quadratic model with random slope effects. FA trajectories are normalized by each tract’s predicted week 1 value to enable simultaneous visualization of all tracts. FA declines more rapidly in the early weeks post-injury before reversing direction, rather than following a simple linear trajectory. (B) FA trajectories over time in three example tracts: forceps major (commissural tract), right anterior thalamic radiation (projection tract), and right superior longitudinal fasciculus (association tract).

To formally test whether the recovery trajectories of each DTI metric followed a curved (rather than straight) pattern over time, we used mixed-effects models that incorporated either a linear or quadratic term for weeks since TBI, assessing how weeks since TBI influenced changes in each DTI metric. Model fit was evaluated and compared between models using Akaike Information Criterion (AIC) and ANOVA model comparisons (Table 1).

**Table 1:**
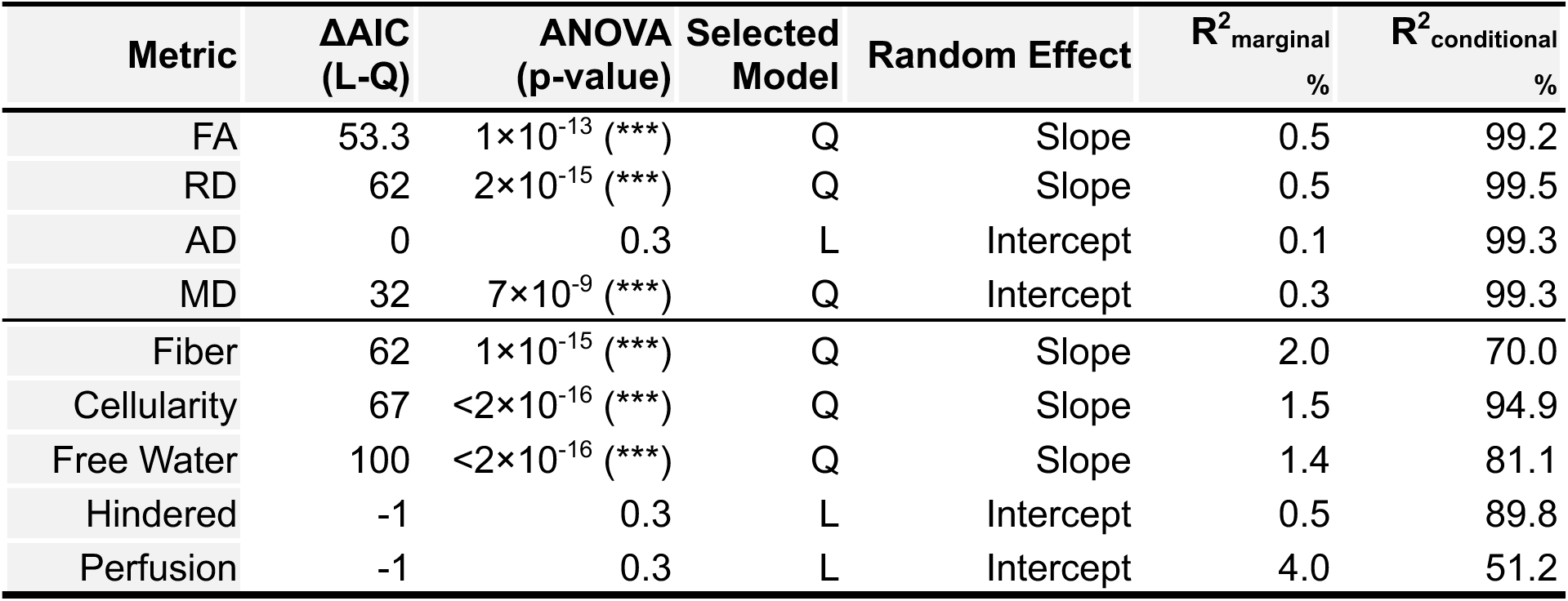
Model Selection across White Matter Metrics in the TBI Patient, comparing linear (L) and quadratic (Q) models using the Akaike Information Criterion (AIC) and ANOVA-based significance testing favoring quadratic models (*p* < 0.001 ***). ΔAIC (L-Q) represents the difference between linear and quadratic models (values >10 favor quadratic models; Burnham & Anderson, 2002). Model selection followed a two-step process: (1). *Model Stability Assessment*: Each model was tested across 1000 bootstrapped simulations to check for convergence and avoid boundary fit issues. (2). *Random Effects Evaluation*: The inclusion of random effects was tested by comparing models with and without them using ANOVA, to determine whether a simpler model without random effects provided a better fit. If a model failed across any simulation or the random effect was non-significant, the model was simplified stepwise (Barr et al., 2013): Quadratic model with random slope → Quadratic model with random intercept → Linear model with random slope → Linear model with random intercept → Simple linear regression (if necessary). The “Selected Model” column shows the best-fitting model, and “Random Effect” indicates whether a slope or intercept was retained as a random effect. R^2^marginal represents the proportion of variance explained only by the fixed effects (i.e., weeks since TBI, head motion, and for quadratic models, weeks since TBI^2^). R^2^conditional represents the total variance explained by both fixed and random effects, incorporating variability due to individual tracts.

| Metric | $\Delta AIC$<br>(L-Q) | ANOVA<br>(p-value) | Selected<br>Model | Random Effect | $R^2_{\text{marginal}}$<br>% | $R^2_{\text{conditional}}$<br>% |
| --- | --- | --- | --- | --- | --- | --- |
| FA | 53.3 | $1 \times 10^{-13}$ (***) | Q | Slope | 0.5 | 99.2 |
| RD | 62 | $2 \times 10^{-15}$ (***) | Q | Slope | 0.5 | 99.5 |
| AD | 0 | 0.3 | L | Intercept | 0.1 | 99.3 |
| MD | 32 | $7 \times 10^{-9}$ (***) | Q | Intercept | 0.3 | 99.3 |
| Fiber | 62 | $1 \times 10^{-15}$ (***) | Q | Slope | 2.0 | 70.0 |
| Cellularity | 67 | $< 2 \times 10^{-16}$ (***) | Q | Slope | 1.5 | 94.9 |
| Free Water | 100 | $< 2 \times 10^{-16}$ (***) | Q | Slope | 1.4 | 81.1 |
| Hindered | -1 | 0.3 | L | Intercept | 0.5 | 89.8 |
| Perfusion | -1 | 0.3 | L | Intercept | 4.0 | 51.2 |

In the TBI patient, quadratic models provided a significantly better fit than linear models for FA, RD, and MD, as indicated by ANOVA model comparisons (ps < 10^-8^) as well as AIC differences (all Δ*AIC*_*Linaer-Quadratic*_ > 10, suggesting minimal support for the linear model; Burnham & Anderson, 2002). Post-hoc modeling revealed that within individual white matter tracts, the quadratic model provided a strong fit for the post-TBI trajectory of FA and RD. On average, the model explained a remarkably large portion of the variance in these measures, with mean R² values across tracts of 0.49 (SD = 0.22) for FA and 0.49 (SD = 0.19) for RD (Fig. 1, S1A, B). Across tracts, 10 out of 18 showed a significant relationship between FA and weeks since TBI², while 9 out of 18 showed a significant relationship for RD. In many tracts, the model accounted for more than 50% of the variance in FA and RD (Fig. 1A, S2, S1A), with particularly strong fits in tracts such as the forceps major, where explained variance exceeded 75% (Fig. 1B, S1B). For MD, the quadratic model explained less variance overall, with a mean R² of 0.30 (SD = 0.19; Fig. S1E, F). Across tracts, 4 out of 18 showed a significant relationship between MD and weeks since TBI². AD, in contrast, was best explained by a linear model, with 6 out of 18 tracts showing a significant but much weaker relationship between AD and weeks since TBI (mean R² = 0.10, SD = 0.17; Fig. S1C, D).

**Figure 2:**
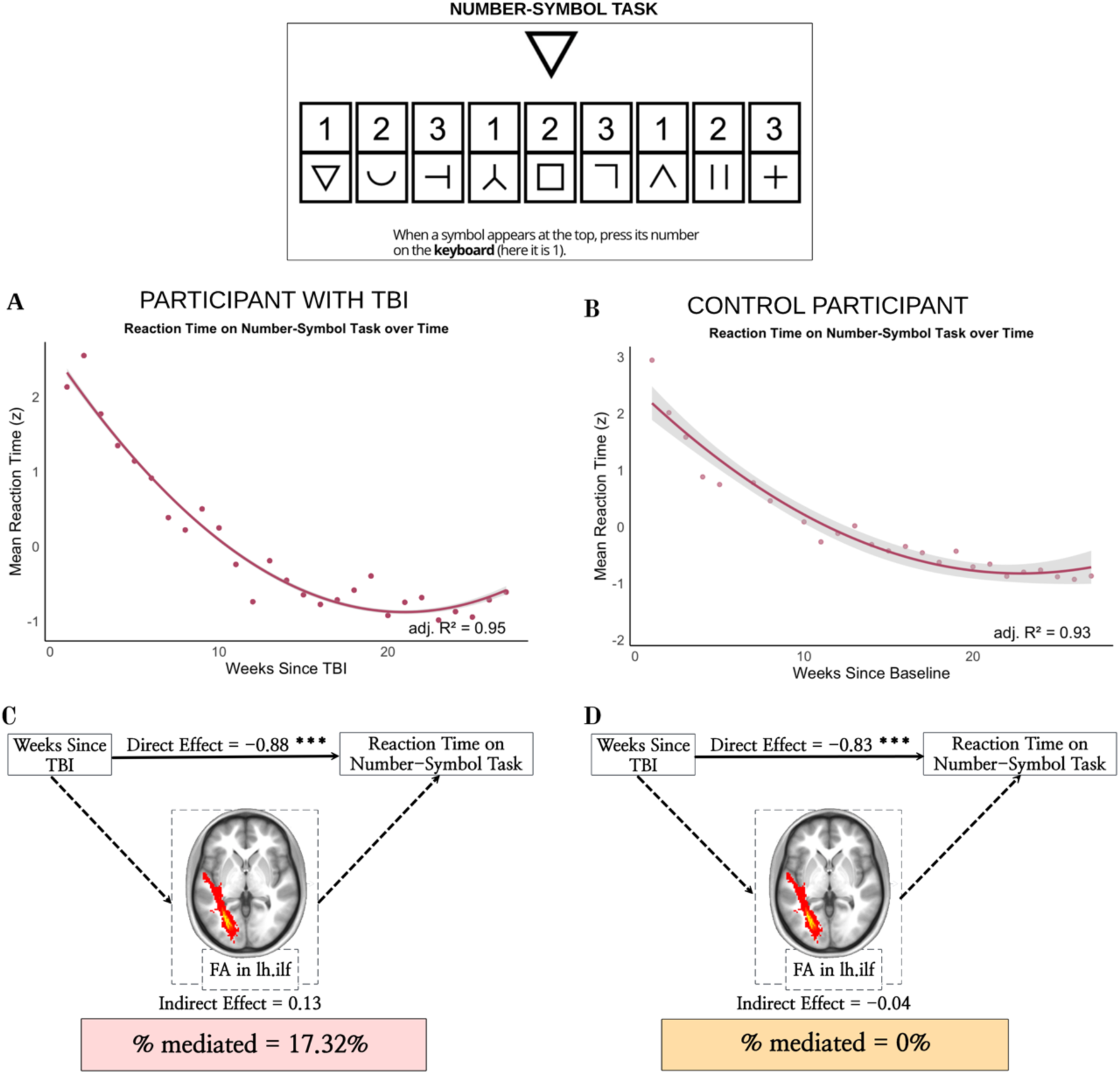
Cognitive Improvements on the Number-Symbol Task Mediated by White Matter-Driven Recovery in TBI. Both the participant with TBI (A) and control participant (B) improved on the number-symbol cognitive task over time, with reaction times decreasing rapidly in the early weeks before stabilizing. To determine whether these improvements reflect practice effects (due to repeated exposure) or longitudinal recovery effects (linked to TBI-related white matter changes), we tested this relationship within a causal mediation framework. In the participant with TBI (C), time since injury strongly predicted faster reaction times (negative direct effect). However, mediation analysis revealed that lower fractional anisotropy (FA) in the left inferior longitudinal fasciculus (ILF) partially enhanced (17%) the beneficial direct effect of weeks since TBI on cognitive improvement. Thus, reductions in FA during recovery contributed to greater cognitive improvement than would be expected by time alone. This mediation effect was completely absent in the control participant (D; mediation effect = 0%), where reaction time improvements occurred independently of white matter changes.

Examining overall recovery patterns, FA initially declined before stabilizing and then increasing over time (Figure 1A, B). In contrast, RD increased in the early weeks post-TBI before gradually decreasing (Fig. S1A, B). MD followed a similar nonlinear trajectory to RD, showing an initial increase followed by a gradual decline (Fig. S1E, F). Unlike these metrics, AD exhibited a more linear trend, generally showing a steady increase over time (Fig. S1C, D).

In the control participant, post-hoc modeling indicated that FA within individual tracts remained relatively stable over time, showing only minor fluctuations compared to the pronounced trajectory observed in the TBI patient (mean R^2^=0.06, SD =0.07; Fig S3A, B). No tracts showed a significant relationship between FA and weeks since TBI². Similarly, RD (mean R^2^=0.03, SD=0.07; Fig S3C, D), and MD (mean R^2^=0.07, SD=0.09, FigS3G, H) in the control participant lacked the distinct increase-and-stabilization pattern observed in the TBI patient. No tracts showed a significant relationship between RD and weeks since TBI² or between MD and weeks since TBI². This observation suggests that variation in DTI measures in an uninjured participant reflect random variability rather than biologically meaningful white matter remodeling.

### 2. Extent of White Matter Restructuring Correlates with Duration of Change

In examining FA and RD trajectories (Fig. 1A, S1B), we observed that tracts exhibiting larger deviations in FA/RD from the initial baseline also seemed to have later inflection points—where the trajectory shifts direction (i.e., FA transitions from decline to increase and RD from increase to decrease). To formally test this relationship, we measured FA and RD changes up to their respective inflection points and correlated them with inflection timing. FA exhibited a significant negative correlation (Fig. S4A, R=-0.67, p=0.003; mean inflection point = 18 weeks, SD=3 weeks), indicating that tracts with later inflection points exhibited greater FA reductions. Conversely, RD exhibited a significant positive correlation (Fig. S4B, R=0.71, p=0.003; mean inflection point = 20 weeks, SD=3 weeks) indicating that tracts with later inflection points exhibited greater RD increases. These results suggest that when the post-TBI white matter remodeling process is more extensive, that process is also more temporally extended.

### 3. White Matter Trajectories Correspond with Recovery of Cognitive and Emotional Processes After TBI

Cognitive and emotional functions are often immediately impacted by TBI but can eventually recover. To assess whether changes in white matter contribute to cognitive and emotional recovery following TBI, we examined whether trajectories of DTI measures mediate improvements in Number-Symbol task performance and reductions in anxiety and depression symptoms over time.

In the TBI patient, we observed that reaction times improved over time (Fig 2A), with a curvilinear trajectory that generally followed the trajectory of DTI changes (Fig 1). However, reaction times on cognitive tasks often improve with repeated administration due to practice effects (Bartels et al., 2010; Holm et al., 2022; Cook, Ramsay, & Fayers, 2004). Indeed, a similar trajectory of improvement was observed in the control participant (Fig 2B).

To determine whether these reaction time improvements depended on white matter changes, we first used a quadratic mediation network to test whether DTI metrics in specific white matter tracts mediated cognitive improvements. To ensure statistical robustness, mediation models were bootstrapped 5000 times. This analysis revealed that for the TBI patient, FA in the left inferior longitudinal fasciculus (ILF) showed a significant indirect effect (*β*_*Indircet*_ = 0.13, p = 0.03), while the direct effect remained negative and significant (*β*_*Dircet*_ = −0.88, p < 0.001), yielding a total effect of *β*_*Total*_ = −0.75 ( p < 0.001). The proportion mediated was -17.3% (p = .03), suggesting that reductions in FA in the left ILF during recovery contributed to greater cognitive improvement than would be expected based on time alone. In the right ILF, both RD and MD showed a significant indirect effect (*β*_*Indircet*_ = −0.18, p = 0.01 for both metrics), alongside a significant direct effect (*β*_*Dircet*_ = −0.56 and *β*_*Dircet*_ = −0.58 respectively, p < 0.001) with total effects of *β*_*Total*_ = −0.75 and *β*_/0+%1_ = −0.76 respectively (p < 0.001).

The proportion mediated was 24.6% (p = .01) for RD and 23.8% (p=0.01) or MD, indicating partial mediation. These findings suggest that structural changes in white matter reflect the trajectory of cognitive recovery.

In contrast, no significant mediation effect was observed in the control participant (Fig. 2D), where reaction time improvements occurred independently of any change in any DTI metric (FA, RD, AD, or MD). Across all tracts, indirect effects ranged from -0.06 to 0.09, with all p-values non-significant (p = 0.13 – 0.98), indicating no evidence of mediation. Similarly, proportion mediated estimates ranged from 0% to 7.5%, with all p-values non-significant (p = 0.13 – 0.98). Although direct effects remained highly significant (β = -0.78 to -0.96, p < 0.001), the absence of a significant indirect effect indicates that reaction time improvements in controls were not mediated by white matter changes.

For comparison, a simpler linear mediation model was also conducted, wherein all DTI metrics and reaction time were modeled as linear functions of weeks since TBI (or weeks since enrollment). This analysis replicated the quadratic mediation findings and identified additional tract-level contributions, including the forceps major as well as the right inferior and superior longitudinal fasciculus, suggesting more widespread white matter involvement in cognitive recovery (Table 2).

**Table 2:**
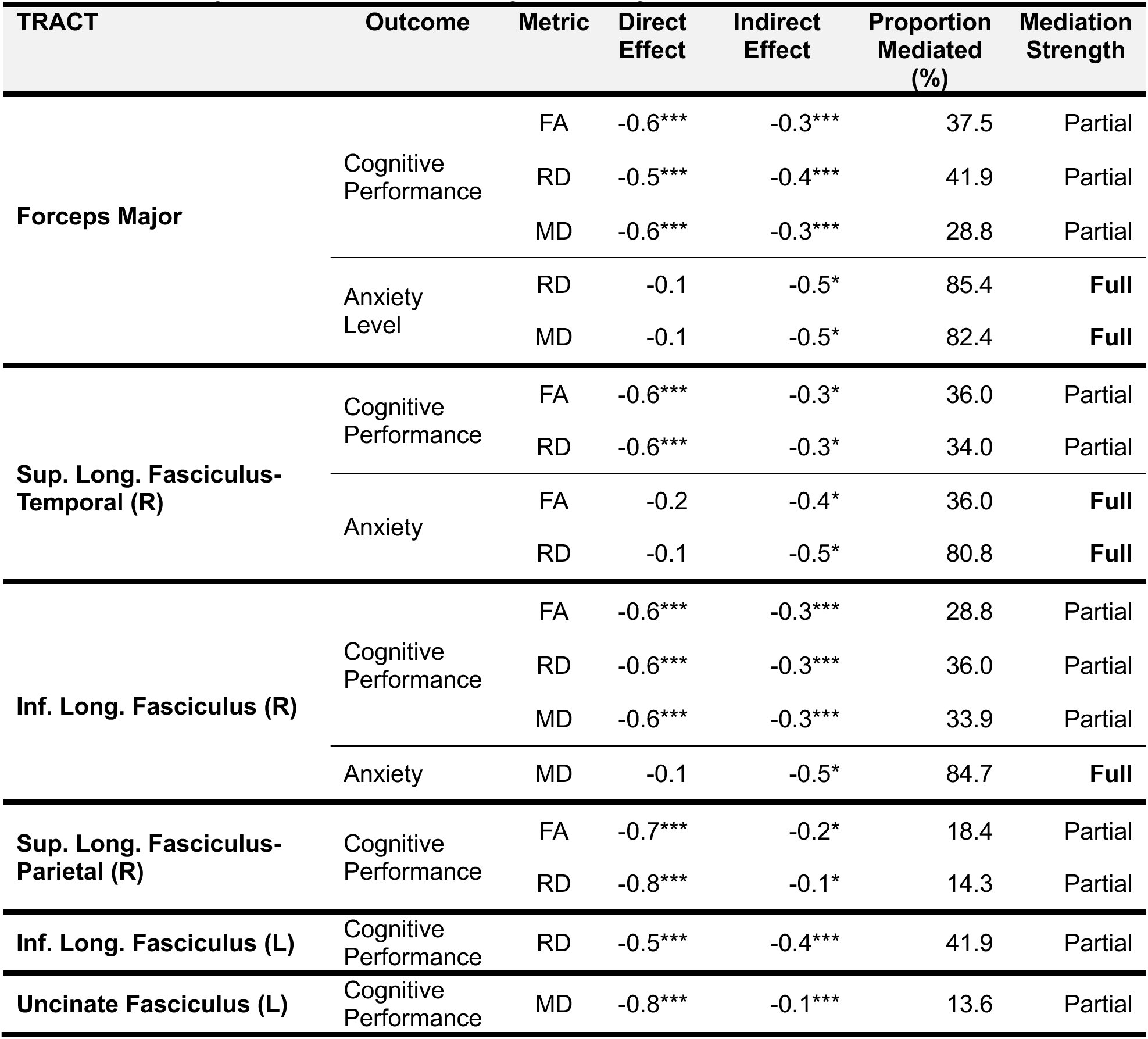
L**i**near **Mediation Models for Reaction Time Performance on Number-Symbol task and Mean Anxiety levels on Beck Anxiety Inventory** Statistically significant results from linear mediation models assessing how DTI metrics mediate cognitive and emotional outcomes post-TBI. The models examine the relationship between weeks since TBI and (1) reaction time performance on the Number-Symbol task (cognitive outcome) and (2) mean anxiety levels (emotional outcome), with mediation effects assessed through fractional anisotropy (FA), radial diffusivity (RD), axial diffusivity (AD), mean diffusivity (MD). No mediation effects for AD were found for any tract or outcome. Mediation effects were categorized as either partial (when indirect effects explained part of the relationship between weeks since TBI and outcome) or full (when indirect effects through the DTI metrics accounted for most of the observed association). Statistical significance is denoted as p<0.05 (*) and p < 0.001(***). FA and RD predominantly mediated cognitive outcomes, with partial mediation observed in the forceps major, right superior longitudinal fasciculus (temporal and parietal projections), bilateral inferior longitudinal fasciculus, and left uncinate fasciculus. RD and MD primarily mediated anxiety levels, with full mediation observed in the forceps major, right superior longitudinal fasciculus (temporal projections), and right inferior longitudinal fasciculus.

Beyond cognitive improvements, we also examined emotional recovery following TBI. Anxiety symptoms showed a sustained reduction, following a similar trajectory to DTI measures, whereas depression symptoms rebounded earlier (Fig. S4). We employed the same mediation approach as above to test for effects on post-TBI recovery of emotional symptoms. Only a linear mediation model yielded significant effects. Notably, the same tracts implicated in cognitive improvements—the forceps major and superior longitudinal fasciculus—also mediated reductions in anxiety symptoms (Table 2), suggesting overlapping white matter mechanisms underlying both cognitive and emotional recovery. FA, RD, and MD in these tracts *fully mediated* reductions in anxiety. Specifically, RD and MD in the forceps major (*β*_*Indircet*_ = -0.48, *p* = .02; *β*_*Indircet*_ = -0.46, *p* = .01, respectively), mediated 85.4% and 82.4% of the total effect (*p* = .03 and *p* = .02). FA and RD in the superior longitudinal fasciculus (*β*_*Indircet*_ = -0.42, *p* = .04; *β*_*Indircet*_= -0.46, *p* = .01, respectively), mediated 72.4% and 80.8% of the total effect (*p* = .04 and *p* = .02). No significant mediation effects were found for depression symptoms. *See Table 2 for full mediation statistics*.

### 4. Characterization of White Matter Recovery Using DBSI: Linking Structural, Inflammatory, and Vascular Mechanisms Post-TBI

To disaggregate the complex microstructural changes following TBI, we leveraged DBSI to capture distinct biological processes—axonal integrity, neuroinflammation, and vascular contributions—that are not fully resolved by DTI (Wang et al., 2019a, 2015, 2011). DBSI metrics can more specifically distinguish the separate biophysical effects on water diffusion caused by fiber loss or demyelination, inflammation, edema, and vascular changes (Wang et al., 2015, 2014, 2011, Chiang et al., 2014). DBSI metrics—including fiber ratio, cellularity ratio, free water ratio, hindered ratio, and perfusion ratio—were extracted from 18 major white matter tracts.

Longitudinal trajectories were modeled over the 27 weeks post-TBI to assess distinct recovery patterns.

As for the DTI metrics, we used mixed-effects models that incorporated either a linear or quadratic term for weeks since TBI, assessing how weeks since TBI influenced changes in each DBSI metric. Model fit was evaluated and compared between models using Akaike Information Criterion (AIC) and ANOVA model comparisons (Table 1).

Quadratic models provided a significantly better fit than linear models for fiber ratio, cellularity ratio, and free water ratio, as indicated by ANOVA model comparisons (ps < 10^-8^) as well as AIC differences (all Δ*AIC*_*Linaer-Quadratic*_ > 10, suggesting minimal support for the linear model (Burnham & Anderson, 2002). In contrast, linear models provided a better fit for hindered ratio and perfusion ratio (ANOVA model comparisons yielding p = 0.25 and p=0.22 respectively, and Δ*AIC*_*Linaer-Quadratic*_ = 1 for both measures).

Post-hoc modeling revealed significant relationships with weeks since TBI² in three of 18 tracts for fiber ratio (mean R² =0.10, SD = 0.16), cellularity ratio (mean R^2^ =0.24, SD=0.17), and free water ratio (mean R^2^ =0.16, SD=0.15). Notably, the right cingulum showed the strongest quadratic effects for fiber ratio (R² = 0.51) and free water ratio (R² = 0.60), while the forceps major exhibited the strongest effect for cellularity ratio (R^2^ = 66%). In contrast, hindered ratio and perfusion ratio were better explained by linear models, with six tracts each showing significant effects of weeks since TBI, but lower explanatory power overall (mean R² for hindered ratio = 0.10, SD = 0.08; mean R² for perfusion ratio = 0.07, SD = 0.08).

The overall patterns of changes followed an inverted U-shaped trajectory for fiber ratio, with early increases giving way to declines at later time points. In contrast, cellularity ratio and free water ratio exhibited a U-shaped course, characterized by initial reductions followed by later increases. Meanwhile, hindered diffusion increased linearly, whereas perfusion ratio declined linearly (Fig 3, Table 1).

**Figure 3.**
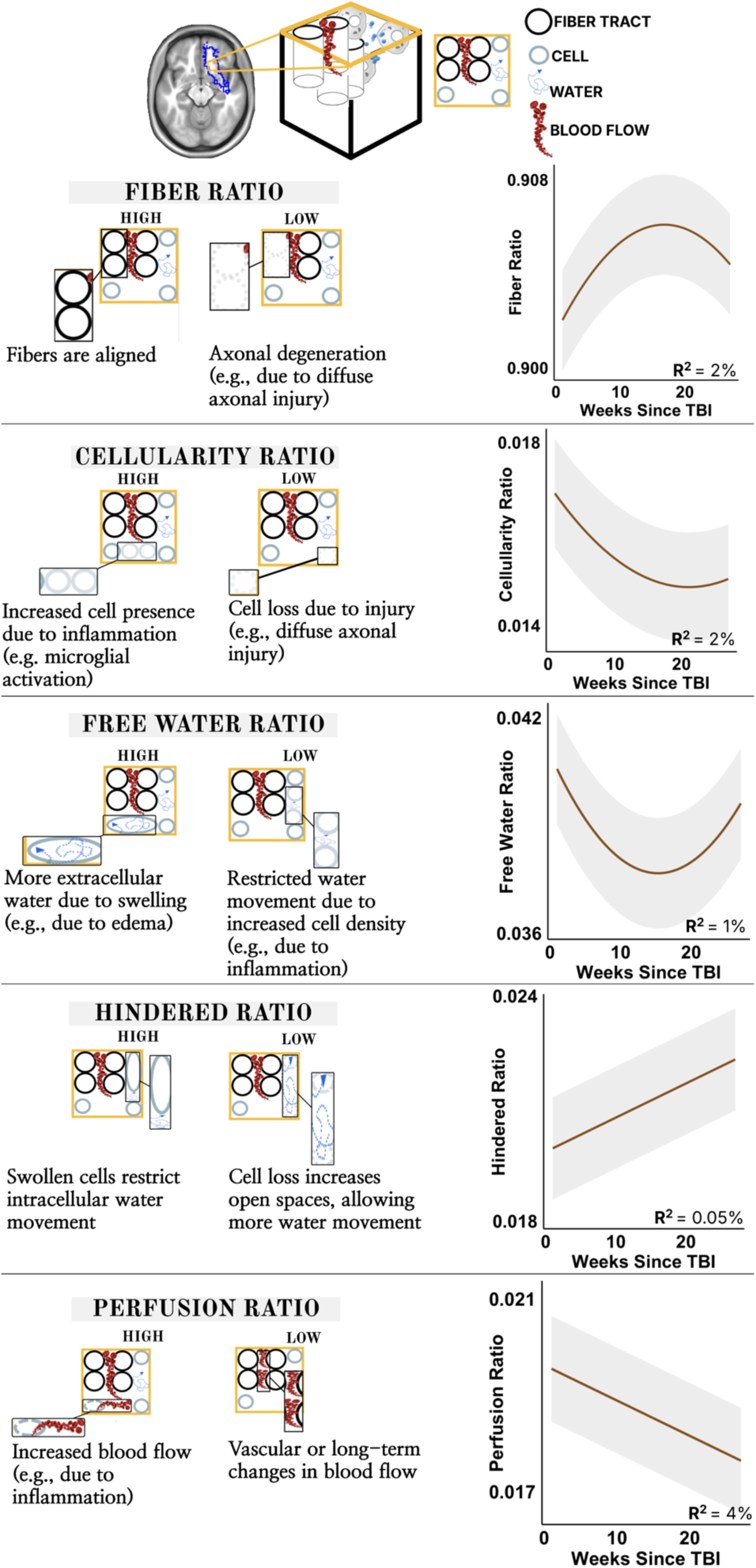
Model Interpretation of DBSI Metrics Following TBI. Top: representation of a white matter tract (outlined in blue) and a single voxel (cube), illustrating how diffusion signals are influenced by multiple biophysical factors at the cellular level, which can be differentiated using Diffusion Basis Spectrum Imaging (DBSI). *Fiber Ratio*: Left—A high Fiber Ratio indicates well-aligned fibers, while a reduction suggests axonal degeneration or demyelination, as observed in diffuse axonal injury. Right—Following TBI, Fiber Ratio initially increases before declining, potentially reflecting a transient prominence of axonal fibers in the diffusion signal before later structural remodeling. *Cellularity Ratio*: Left—An increase in cellularity corresponds to immune cell infiltration, indicative of an inflammatory response, while a decrease suggests cell loss. Right—Cellularity initially declines post-TBI, likely due to early cell loss, before rebounding over time, suggesting neuroinflammatory responses and tissue remodeling. *Free Water Ratio*: Left— Extracellular free water levels increase with edema and decrease with inflammation-driven cell accumulation. Right—The Free Water Ratio follows a U-shaped trajectory over time, initially decreasing before increasing again, likely due to evolving edema and tissue remodeling. *Hindered Water Ratio*: Left—Intracellular water movement is restricted by swollen cells, while cell loss increases open extracellular space, facilitating greater water diffusion. Right—The Hindered Water Ratio shows a small increase over time, suggesting progressive microstructural alterations that facilitate more intracellular water movement. *Perfusion Ratio*: Left—Elevated perfusion suggests increased blood flow due to inflammation, whereas a reduction may indicate vascular remodeling or long-term changes in cerebral blood flow. Right—The Perfusion Ratio declines progressively post-TBI, indicating potential reductions in inflammation-driven hyperemia or long-term microvascular alterations. Each graph presents longitudinal trajectories of these DBSI-derived metrics across weeks post-TBI, with shaded regions representing confidence intervals. In each graph, R^2^ values indicate marginal effects, i.e., the proportion of variance explained only by the fixed effects (i.e., weeks since TBI, head motion, and for quadratic models, weeks since TBI^2^).

### 5. Non-linear Recovery of FA and RD at 1-year follow-up

To evaluate the long-term trajectory of white matter recovery post-TBI, we analyzed all DTI metrics at a 1-year follow-up. The previously observed quadratic patterns persisted out to 1 year. FA, which initially declined before stabilizing, exhibited recovery to nearly the initial post-injury levels (Fig 4A). Similarly, RD increased before stabilizing and returned close to initial levels (Fig 4B), while MD followed a parallel trajectory (Fig 4D), reinforcing a shared nonlinear recovery process. In contrast, AD exhibited greater variability across tracts, with a weak linear fit suggesting inconsistent associations with recovery (Fig 4C), making it a less robust marker than FA or RD.

**Figure 4:**
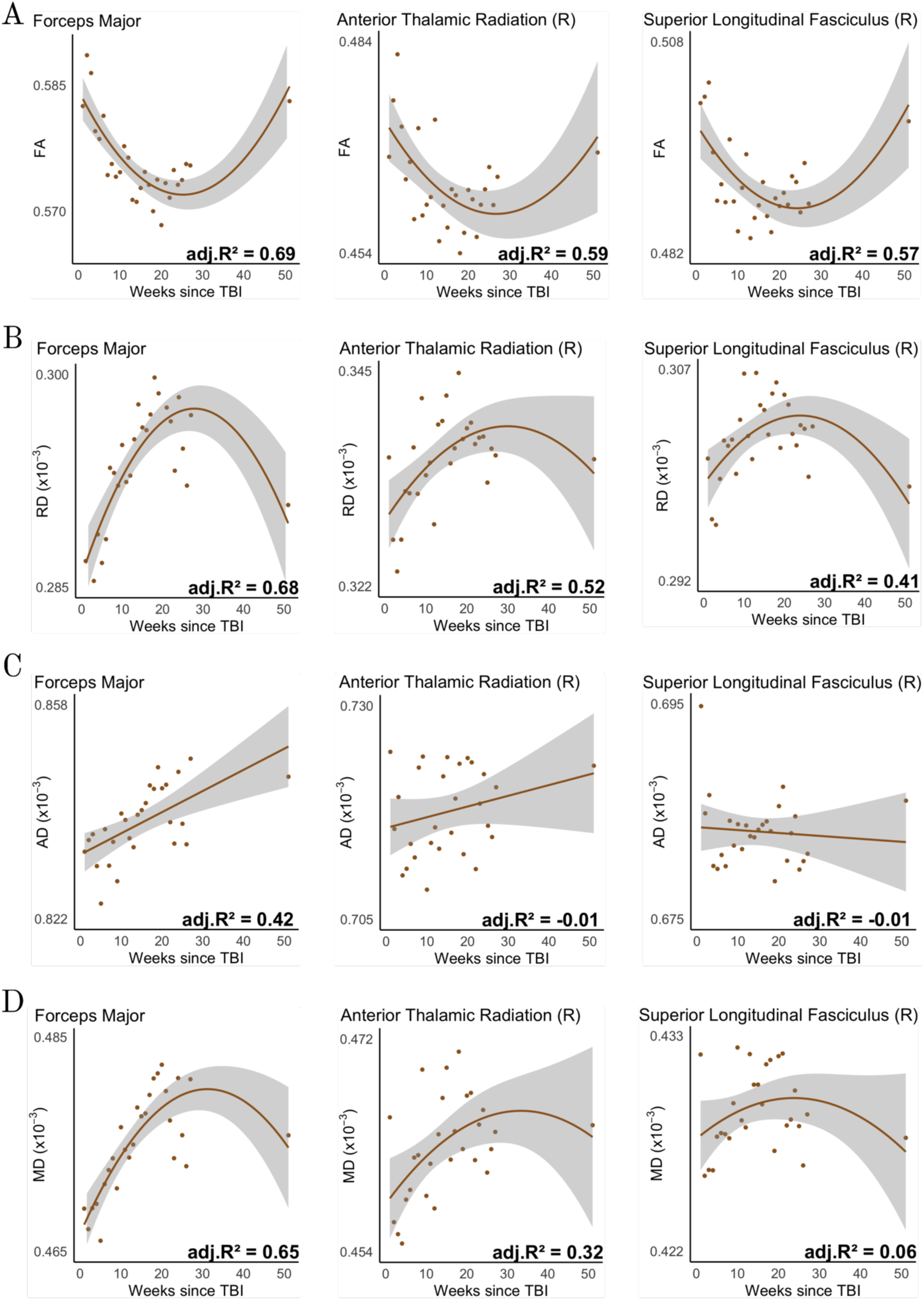
**Persistence of Non-Linear Trajectories of DTI Metrics up to One-Year Post-TBI Across Specific White Matter Tracts**: Each row corresponds to a specific DTI metric, tracking its trajectory over one-year post-TBI. The shaded regions represent 95% confidence intervals, while adjusted R² values (adj. R²) quantify the proportion of variance explained by the model. (A) Fractional Anisotropy (FA), initially declined post-injury but demonstrated a quadratic recovery trajectory, approaching early post-injury levels by one year. The high adj. R² values across tracts indicate a strong fit for the quadratic model, reinforcing the nonlinear nature of FA recovery. (B) Radial Diffusivity (RD) exhibited an early post-injury increase followed by stabilization and a gradual return toward initial levels. (C) Axial Diffusivity (AD) showed a more variable trajectory, with some tracts exhibiting a subtle increase over time. However, low adj. R² values suggest weak or inconsistent associations with post-injury recovery, implying that AD may not be as robust of a marker for TBI recovery as FA or RD. (D) Mean Diffusivity (MD), exhibited a similar quadratic pattern to RD, with an initial increase followed by stabilization and partial reversal.

Across all tracts (Fig. S6), right-hemisphere tracts adhered more consistently to the quadratic model, whereas left-sided tracts showed greater deviations over time. This pattern may be influenced by the specific injury profile of the TBI patient, who sustained a right temporal subdural hematoma.

## Discussion

This densely sampled, precision longitudinal study aimed to elucidate the individual-level trajectory of white matter remodeling following traumatic brain injury. We found that DTI measures of white matter restructuring—particularly changes in FA and RD—follow a non-linear, parabolic trajectory over the first year (Fig 1, S1, 4).

Complementary DBSI-analyses suggested early cell loss followed by inflammatory processes leading to persistent alterations in white matter microstructure, such as long-term microvascular changes (Fig 3). These changes are not mere snapshots of damage; rather, they have direct functional consequences, as changes in fiber integrity measures are associated with the rate of cognitive recovery and anxiety symptoms (Fig 2, Table 2).

Despite increasing recognition of white matter plasticity post-TBI, the interpretation of recovery trajectories remains hindered by the rigid and inconsistent operationalization of subacute (> 1 week to ≤ 3months) and chronic phases (>3months; Wallace, Mathias, & Ward, 2018). The absence of clear differentiation in the underlying neurobiological processes and recovery dynamics across these intervals raises the question of whether these categories are biologically meaningful or merely artifacts of inconsistent definitions. Rather than adhering to fixed time windows, our findings suggest that white matter recovery is neither staged nor linear, but follows a dynamic trajectory, marked by distinct inflection points (Fig 1, S1, S2). This perspective shifts the focus from predefined temporal phases to transitional periods, where inflection points mark key shifts in the recovery process. Indeed, the timing of these transitions correlates with the extent of white matter remodeling required (Fig S2), suggesting that recovery phases are not dictated by fixed post-injury intervals, by the degree of structural alterations and the time needed for tract-specific reorganization. By identifying inflection points in white matter remodeling, we may establish a more precise and individualized framework for assessing injury progression and guiding treatment.

These findings challenge the traditional view that reduced FA in TBI simply reflects white matter damage. FA has long been considered a marker of “white matter integrity” (Leow et al., 2009, Shenton et al., 2012, Kochunov et al., 2012, 2007, Kraus et al., 2007, Abdullah et al., 2022), with lower FA in TBI patients compared to healthy controls often interpreted as evidence of structural deterioration (Wallace, Mathias, & Ward, 2018, Hulkower et al., 2013). However, our results show that FA declines before shifting direction over a longer period than expected if it only reflected immediate or irreversible damage. If FA were solely a marker of structural deterioration, its initial magnitude of change would scale with the severity of the insult, but not necessarily influence how long these changes persist. Within this framework, more affected tracts would be expected to show steeper FA reductions early on, but without considering other microstructural processes at play, these changes would be expected to either plateau or continue declining monotonically (e.g., due to Wallerian degeneration; Bendlin et al., 2008, Edlow et al., 2016, Harris et al., 2016, Lindsey et al., 2023). Instead, we observed that tracts exhibiting larger FA changes also showed later inflection points (Fig S2), suggesting that white matter alterations unfold over an extended period, following a trajectory that is not easily explained by passive degeneration alone. More strikingly, rather than signaling ongoing deterioration, these extended FA changes were associated with improved cognitive and emotional outcomes (Fig 2, S5, Table 2). This observation is difficult to reconcile with the idea that FA simply reflects a breakdown in structural integrity. Instead, our findings suggest that FA may also capture dynamic, adaptive processes that influence long-term structural changes.

Ultimately, the conventional framework for interpreting FA in TBI appears incomplete. These findings underscore the need for a more nuanced understanding of FA’s role in brain injury, one that accounts for its evolving relationship with functional recovery over time.

An improved interpretation of FA changes is enabled by our complimentary use of DBSI measures. A fundamental challenge in interpreting DTI metrics is that measures like FA can be influenced by multiple physiological processes (Figley et al., 2022). FA stabilization might mask ongoing axonal repair or degeneration (Pierpaoli et al., 2001). In our study, we observed an initial decline in FA immediately following TBI, accompanied by reductions in both the cellularity ratio and free water ratio (Fig 3C, D). The decrease in cellularity suggests significant cell loss following TBI, while the lower free water ratio indicates a lack of early extracellular water accumulation—likely due to tissue collapse following cell loss (Wang et al. 2019a, 2011, Chiang et al. 2014). Under normal conditions, cellular swelling in response to stress would restrict water movement and increase hindered diffusion. However, in this early phase, cell loss appears to outweigh swelling, leading to disrupted tissue architecture, which may explain the observed reduction in FA (Fig 1; Wang et al. 2019a, 2011, Chiang 2014). Concurrently, with fewer non-axonal cells present, the remaining intact, well-organized axons may become more prominent in the diffusion signal, resulting in a transient rise in the fiber ratio (Chiang et al., 2014, Wang et al., 2017).

As the injury evolves, inflammatory processes likely contribute to structural remodeling. Immune cell infiltration and edema lead to increases in both the cellularity and free water ratios and a corresponding decrease in hindered ratio (Fig 3E; Wang et al., 2011). At this stage, the tissue may be undergoing structural remodeling. These processes may improve fiber alignment among surviving axons or represent reorganization that partially compensates for earlier damage. While excess fluid from edema may initially disrupt fiber structure, the resolution of inflammation and edema may allow for partial recovery of fiber integrity, potentially contributing to the observed shift in FA trajectory (Wang et al. 2019a, 2011, Chiang et al. 2014). However, this process does not appear to reach a stable endpoint, as the perfusion ratio continues to increase throughout the observed time frame (Fig 3F). Persistent perfusion changes suggest ongoing metabolic activity, potentially linked to sustained remodeling (Wang et al., 2019b) rather than a return to a pre-injury baseline. Thus, white matter recovery is not a simple return to baseline, but a dynamic, evolving process shaped by fluid shifts, immune responses, and prolonged vascular adaptations.

By studying the process of recovery from TBI within rather than across individuals, the present work may more clearly identify biological mechanisms linked with recovery from TBI—a fundamentally within-individual process—than classic cross-individual, group-average studies. There is growing recognition that results obtained from intra-individual human studies may not be appropriate to infer effects occurring within individuals (Fisher et al. 2018). This so-called “ecological fallacy” (Robinson, 2009, Schwartz, 1994) is a particular concern in human neuroscience (Cragg et al., 2019) and results from issues such as Simpson’s paradox (Blyth, 1972, Kievit et al., 2013, Wagner, 1982), in which within-individual effects can be masked or even appear to reverse direction in cross-individual analyses (see Mattoni et al., 2025 for further discussion). Given the expected cross-individual spatial variability of injury and nonlinear temporal dynamics of recovery, TBI may be a condition in which the ecological fallacy is particularly problematic. Thus, precision imaging approaches studying within-individual effects, such as the one employed here, will be needed for effective clinical translation of DTI measures in TBI (Gratton et al., 2020; Kraus et al., 2023; Laumann et al., 2023; Mattoni et al., 2025).

This work provides a new perspective on post-TBI recovery processes focused on individuals rather than on group averages. The clinical implications of this shift in perspective are significant. If white matter microstructure exhibits distinct periods of decline and reorganization within individual patients, interventions could be timed to coincide with optimal windows for neuroplastic adaptation. For instance, if an inflection point signals the onset of compensatory reorganization, targeted cognitive or rehabilitative therapies may be most effective when aligned with the broader trajectory of structural remodeling, rather than applied uniformly across a broad “chronic” window. Future research should focus on defining these biologically driven intervention windows, allowing for personalized rehabilitation strategies informed by empirical recovery trajectories. Additionally, our findings suggest that persistent inflammatory activity may contribute to prolonged structural alterations, suggesting that TBI is not simply the residual effect of an initial injury, but an evolving pathophysiological process. This distinction has critical implications for late-stage interventions, including anti-inflammatory strategies and neuroprotective therapies, which may help mitigate long-term neurodegeneration and associated cognitive decline or mental health conditions.

Ultimately, our integrated approach—combining dense longitudinal sampling, advanced mixed-model analyses, and the complementary use of DBSI—provides enhanced specificity in interpreting white matter changes following traumatic brain injury (TBI). This strategy not only overcomes many of the traditional pitfalls associated with DTI metrics (Figley et al., 2022) but also lays the groundwork for identifying distinct windows of neuroplastic adaptation. By elucidating tract-specific recovery trajectories, our findings have important clinical implications, suggesting that personalized, time-sensitive interventions may be developed to optimize cognitive and emotional recovery in TBI patients.

## Methods

### 1. Participant and Study Characteristics

The participant with TBI identified as a 21y.o. female with right temporal subdural hematoma (11x8mm) from a motor vehicle collision (MVC; Glascow Coma Scale=15; Teasdale & Jennett, 1974). On arrival at the hospital, she reported a headache but denied nausea, vomiting, new weakness, numbness, or tingling in her extremities. The control participant identified as a 24y.o. male with no history of neurological or psychiatric conditions.

Both participants underwent weekly MRI scans and behavioral assessments for 27 weeks (∼6 months), beginning two weeks after hospital admission for the participant with TBI, followed by an MRI scan and behavioral assessment at their 1-year timepoint. All assessments took place at the Washington University School of Medicine, with Institutional Review Board approval. Both participants provided informed consent for all aspects of the study.

### 2. Magnetic Resonance Imaging (MRI) Acquisition

Neuroimaging data were acquired using a 3T Siemens Prisma scanner with a 64- channel head coil. Each 45-minute session included the acquisition of 4 types of images: (1) a T1-weighted MP-RAGE (sagittal, 224 slices, TE = 3.74 ms, TR = 2400 ms, flip angle = 8°, 0.8 mm isotropic); (2) a T2-weighted image (sagittal, 224 slices, TE = 479 ms, TR = 3200 ms, flip angle = 120°, 0.8 mm isotropic); (3) a field-map scan (three spin-echo image pairs) for dMRI distortion correction, and (4) a diffusion-weighted imaging (DWI) scan (single-shot echo planar, axial, 75 slices, TE = 83 ms, TR = 3500 ms, 2 mm isotropic, four shells: b-values 250, 500, 1000, and 1500 s/mm², 96 encoding directions). Real-time monitoring was implemented using the Framewise Integrated Real-Time MRI Monitoring (FIRMM) system (Dosenbach et al., 2017), which provided immediate motion feedback to minimize artifacts. All sessions were included based on FIRMM-generated motion quality scores.

### 3. Preprocessing Pipeline

#### 3.1. T1 & T2-Weighted Images

Workflow: *FSL FAST → Cross Registration → Mean Image Generation → Atlas Registration → FreeSurfer Surface Reconstruction*.

For each subject, FSL FAST was applied to each of the T1- and T2-weighted images to standardize tissue contrast and segmentation (Zhang et al., 2001). All 27 T1 images were linearly cross registered to each other and averaged across sessions to create a mean T1 image using 4dfp tools (https://readthedocs.org/projects/4dfp/); this procedure was then repeated for the T2 images. The mean T1 image was then linearly registered to MNI atlas space using 4dfp tools. The mean T2 image was linearly registered to the mean T1 image and through to MNI space. Finally, both the mean T1- weighted image and the mean T2-weighted image were processed through the FreeSurfer pipeline (version 7.0) to parcellate cortex, subcortical structures, and white matter.

#### 3.2. Diffusion-Weighted Images (DWI)

Workflow: *Eddy Correction & Topup → Motion Correction → DTI Measure Computation → Atlas Registration* For each session FSL’s Eddy correction and Topup was first applied to the DWI scan, using that session’s field map scans, to correct for distortions caused by eddy currents and MR field distortions (Smith et al., 2004; Andersson & Sotiropoulos, 2016). Second, volumes with framewise displacement >0.5 mm were removed (Baum et al., 2018). Third, the undistorted B0 volume was extracted and linearly registered to the individual’s mean T2-weighted image, and through to MNI space using the T2->MNI registration computed above. Finally, voxelwise diffusion tensors were modeled and DTI metrics were computed from the session using FSL’s DTIFIT. These metrics included (1) *Fractional Anisotropy* (FA), estimated by the extent of directional (anisotropic) water diffusion in each voxel; (2) *Axial Diffusivity* (AD), estimated by the extent of water diffusion along the primary (longest) tensor axis of each voxel; (3) Radial Diffusivity (RD), estimated by the extent of water diffusion perpendicular to the primary axis each voxel; and (4) Mean Diffusivity (MD), estimated as the average water diffusion in all axes of each voxel. These metric maps, as well as the original distortion-corrected DWI images, were registered to MNI space using the DWI->MNI registration.

### 4. Processing Pipeline

#### 4.1. Fiber Tract Definition and DTI Metric extraction

Workflow: *Estimate Tract Probabilities → Integrate with Participant’s DWI Data → Model Fiber Orientations (bedpostX) → Reconstruct Tracts in Participant’s Native Space → Register to MNI space → Extract FA, RD, MD, and AD Measures* First, prior probabilities were estimated by combining a white matter atlas with FreeSurfer-derived cortical and subcortical segmentations (Section 3.1.), determining the likelihood of each tract passing through or near specific anatomical regions. Second, these priors were combined with the participant’s MNI-space diffusion-weighted data to refine tract estimates. Third, FSL’s bedpostX tool was used to fit a ball-and-stick diffusion model, allowing for the representation of up to two distinct fiber orientations per voxel. Fourth, using the estimated fiber orientation vectors and anatomical priors, the TRACULA tool (TRActs Constrained by UnderLying Anatomy; Yendiki et al., 2011) reconstructed each white matter tract, preserving individual anatomical variation and ensuring longitudinal consistency. The final output consisted of probabilistic tract maps representing the most likely trajectories of each of 18 major white matter pathways. These pathways included forceps major, forceps minor, left and right corticospinal tract (CST), inferior longitudinal fasciculus (ILF), uncinate fasciculus (UNC), anterior thalamic radiation (ATR), cingulum-cingulate gyrus bundle (CCG), cingulum-angular bundle (CAB), and superior longitudinal fasciculus (SLF; temporal and parietal branches). Fifth, a streamline threshold of three was applied to the tract maps, to retain only voxels with sufficient probabilistic support for belonging to a given tract. This thresholding process produced individual-specific MNI-space masks for each tract. Finally, we computed mean diffusion metrics (FA, AD, RD, and MD) within each tract mask for every imaging session.

#### 4.2. Diffusion Based Spectrum Imaging (DBSI; Wang et al., 2015, 2014, 2011)

Workflow: *Raw DWI Signal → Multi-Compartment Model Fitting → Regularized Non-Negative Inversion → Extract Diffusion Metrics (Fiber Ratio, Cellularity Ratio, Free Water Ratio, Hindered Ratio, and Perfusion Ratio)* First, the direction-specific diffusion MRI signals within each voxel of the undistorted MNI-space DWI data was decomposed using a multi-compartment model that accounts for different cerebral anatomical and pathological components, including anisotropic white and gray matter fibers, inflammatory cells, extracellular water (edema), and CSF. Second, a regularized nonnegative least-squares analysis was applied to prevent overfitting, incorporating prior knowledge about signal intensities and energy constraints. Third, the model estimated key diffusion parameters, including anisotropic signal fractions, which reflect white matter integrity, axial and radial diffusivity, which assess water movement along and across white matter fibers to infer microstructural health, and an isotropic diffusion spectrum, which quantifies cellular and extracellular properties. For tract-specific extraction, thresholded white matter masks from TRACULA’s MNI-registered tract maps (Section 4.1) were applied to the DBSI-derived parameter maps. This step ensured that diffusion measures were computed only from voxels with a high probability of belonging to a given white matter tract, reducing contamination from surrounding tissue or CSF. Subsequently, mean DBSI metrics were extracted for each tract at every imaging session: (1) *fiber ratio*, quantifying the proportion of intact white matter fibers, (2) *cellularity ratio*, quantifying the extent of restricted diffusion linked to immune cell infiltration, (3) *free water ratio*, quantifying extracellular water content indicative of edema, (4) *hindered ratio*, quantifying intracellular water changes due to swelling or cell infiltration, and, (5) *perfusion ratio*, quantifying vascular blood flow, which may vary with inflammation.

### 5. Cognitive and Emotion Assessments

#### 5.1. Number-Symbol Coding Task

This task simplifies the traditional Symbol Digit Modalities Test (SDMT; Smith 1973; Jaeger 2018) by reducing the number of symbol-digit pairs from nine to three, where each digit (1, 2, or 3) is associated with three different symbols (see Figure 2). Participants are required to identify the correct digit for a given target symbol, providing a measure of processing speed, pattern recognition, and cognitive flexibility while minimizing cognitive load. A digitized version of the task was used in this study, and prior research has demonstrated high correlations between the digital and traditional paper-and-pencil versions (Pham et al., 2021). This task has shown high sensitivity to detecting changes following neurological injury (Jaeger et al., 2018, Galvin et al., 2020), making it a valuable tool for longitudinal cognitive assessment. Moreover, its construct validity ensures that performance is not confounded by anxiety or depression (Jaeger et al., 2018; Table 2), allowing it to be reliably interpreted as a measure of cognitive recovery following TBI.

#### 5.2. **Beck Anxiety Inventory** (**BAI**; Beck et al., 1993)

The BAI is a 21-item self-report measure designed to assess the severity of anxiety symptoms over the past week. Participants rate the presence of physical symptoms (e.g., *“Numbness or tingling”*) and cognitive symptoms (e.g., *“Fear of losing control”*) on a 4-point Likert scale, with higher scores indicating greater symptom severity. The BAI demonstrates high specificity for anxiety symptoms and effectively differentiates anxiety from depressive symptomatology.

#### 5.3. Beck Depression Inventory (BDI; Beck et al., 1996)

The BDI is a 21-item self-report measure designed to assess the severity of depressive symptoms over the past two weeks (modified to a one-week timeframe for this study). Participants rate the presence of physical symptoms (e.g., *“Tiredness or fatigue”*), affective symptoms (e.g., *“Sadness”*), behavioral symptoms (e.g., *“Loss of interest in activities”*), and cognitive symptoms (e.g., *“Self-criticism”*) on a 4-point Likert scale, with higher scores reflecting greater symptom severity. The BDI demonstrates high specificity for depressive symptoms and effectively distinguishes depression from anxiety-related symptomatology.

### 6. Analyses

All statistical analyses were conducted in R (4.4.1, studio version:2024.12.0+467).

#### 6.1. Modeling Trajectories of DTI and DBSI Metrics over Time

To characterize white matter changes over the 27 imaging sessions, linear and quadratic mixed-effects models were fitted to individual DTI and DBSI metrics for each participant to capture potential nonlinear recovery patterns. This modeling framework allowed us to capture general patterns of change in each DTI/DBSI metric over time (e.g., evaluating how weeks since TBI^2^ relates to FA values across all brain tracts; fixed effect), while also accounting for the possibility that different brain tracts have different starting values (i.e., random effect of tract-intercept) and distinct rates of change over time (i.e., random effect of slope; R package, lme4; Bates et al., 2015).

Following best practices in confirmatory hypothesis testing (Barr et al., 2013), we specified a maximal random effects structure, incorporating the most complex model justified by the data to account for variability and enhance generalizability (i.e., starting out with a model that incorporated both random intercept and for tract variation over time). These models were defined as follows:

Linear model

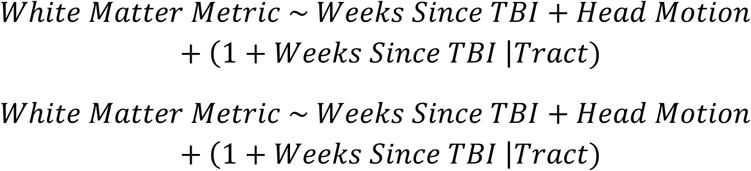

Quadratic model

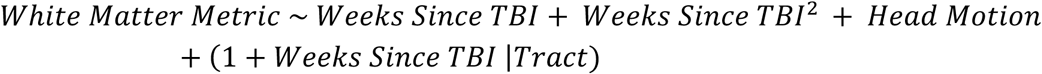

where:

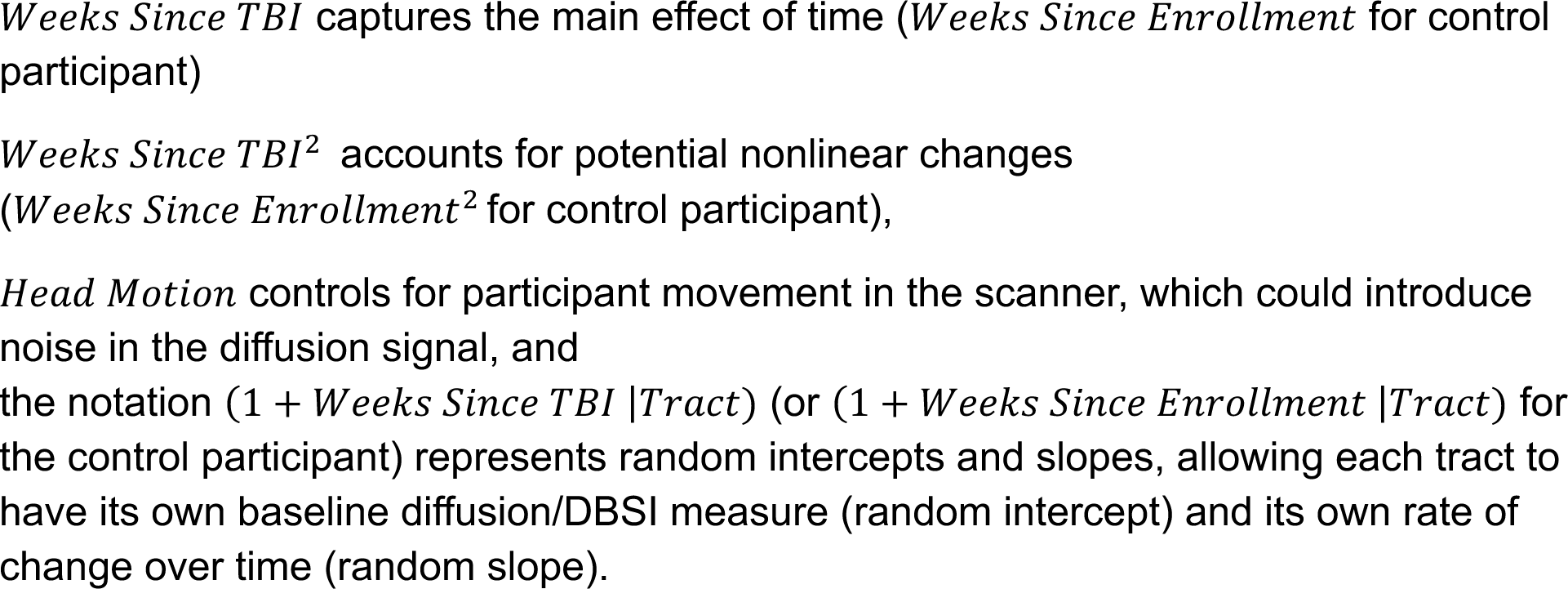

Model fit was evaluated using Akaike Information Criterion (AIC) and ANOVA model comparisons (Table 1). To ensure model stability, all analyses were bootstrapped 1000 times, and models were only accepted if they converged across all iterations. We parsed the separate contributions of fixed effects (R^2^marginal) and combined fixed and random effects (R^2^conditional) using the MumIn package in R (Bartoń K, 2024; Table 1).

Results indicated that FA and RD followed a quadratic trajectory over time (i.e., modeling weeks since TBI^2^ as a fixed effect) with random intercept and slope for tracts (random effects), while MD followed a quadratic trajectory with only random intercept for tracts. In contrast to these quadratic fits, AD best fit a linear mixed-effects model (fixed effect of weeks since TBI) with only random intercept for tracts, Table 1; Result [1]). Model fits for DBSI measures are summarized in Table 1 (Result [4]).

To map these effects onto individual tracts, we estimated adjusted R² values for each tract, modeling tract-wise quadratic effects for FA, RD, and MD and tract-wise linear effects for AD. These R² values were projected onto MNI-registered tract maps (Result [1]).

Linear model (tract-wise)

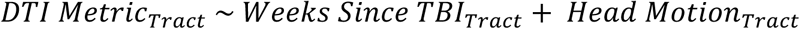

Quadratic model (tract-wise)

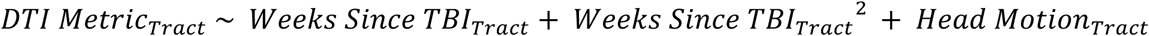

These models were extended to project recovery trajectories up to the one-year timepoint, providing a broader view of long-term white matter changes (Result [4]).

#### 6.2. Modeling Trajectories of Cognitive and Emotion Outcomes and Contributions of White Matter

We modeled the trajectories of cognitive performance (reaction time on accurate trials of the Number-Symbol Coding task) and mean scores on the BAI (anxiety) and BDI (depression) over time for both participants (Result [3]).

To determine whether DTI changes mediated cognitive recovery over time, we implemented a quadratic mediation framework (Preacher & Hayes, 2008). This approach tests whether DTI metrics (FA, RD, MD, AD) mediate improvements in cognitive task performance over weeks since TBI (or weeks since enrollment for the control participant). For each participant and tract, we specifically modeled the following:

Mediator Path Model

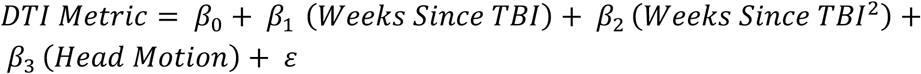

This model estimates how weeks since TBI (or weeks since enrollment for the control participant) affects the DTI metric (the mediator), with *β*_5_ capturing this effect.

Outcome Path Model

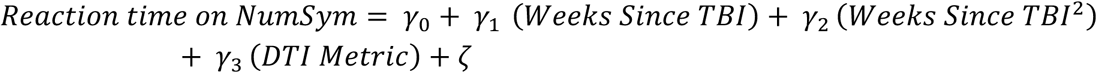

This model tests whether variation in DTI metrics predicts cognitive processing speed —as indexed by reaction time on the Number-Symbol Coding task—while controlling for time since TBI (or enrollment). The coefficient *γ*_6_ captures the effect of DTI changes on reaction time, while *γ*_5_ and *γ*_2_ account for the linear and quadratic effects of weeks since TBI (or enrollment), respectively.

Indirect & Total Effects:

The indirect effect, which quantifies mediation, is computed as:

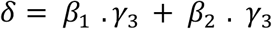

This equation captures the mediation of both the linear and quadratic components of weeks since TBI (or enrollment) on reaction time via changes in DTI metrics.

The total effect of weeks since TBI (or enrollment) on cognitive performance is computed as:

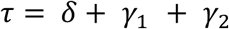

where:

τ represents the total effect of time since injury (or enrollment) on cognition,

δ is the indirect effect mediated by DTI changes, and,

*γ*_1_ *γ*_2_ are the direct unmediated effects of time on reaction time, capturing both the linear and quadratic influence.

Mediation effects were estimated using bootstrapped confidence intervals (5000 simulations; Preacher & Hayes, 2008) to account for non-normality. Proportion mediated was computed as ^7^ x 100, indicating the percentage of the total effect of time mediated by white matter changes. Mediation effects were classified as full, partial, or none based on the significance of indirect and direct effects (Table 2).

This approach addresses limitations of traditional mediation models (Baron & Kenny, 1986) by allowing for causal inference under weaker assumptions. The Preacher & Hayes framework, implemented via the mediation package in R (Tingley et al., 2014), explicitly quantifies indirect and direct effects, provides robust uncertainty estimates, and accommodates nonlinear relationships, making it particularly well-suited for modeling complex neurobiological recovery trajectories.

For comparison, a simpler linear mediation model was also conducted (Table 2), wherein all DTI metrics and reaction time were modeled as linear functions of weeks since TBI (or weeks since enrollment). In this instance:

Mediator Path Model

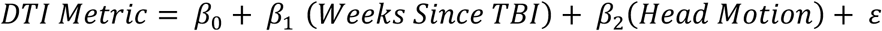

Outcome Path Model

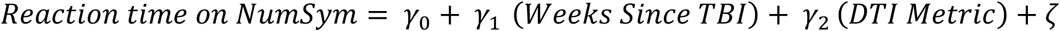

Indirect & Total Effects:

The indirect effect is computed as:

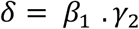

The total effect of weeks since TBI on cognitive performance is computed as:

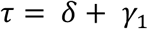

where:

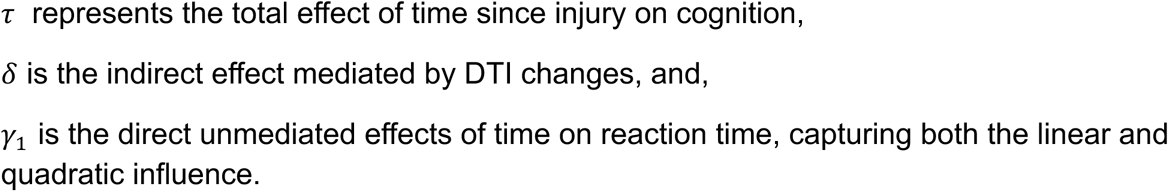

Both quadratic and linear mediation models were replicated for emotional outcomes, with the same mediation path models and indirect and total effects calculations (Table 2). The outcome path model was defined as:

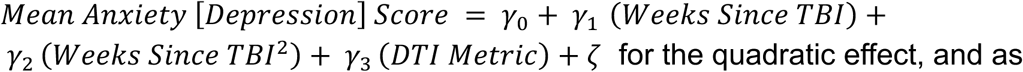

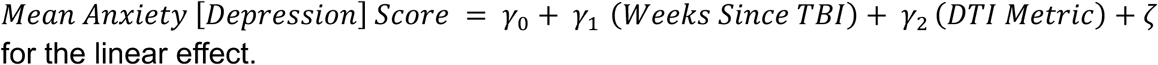

---

**Figure S1:**
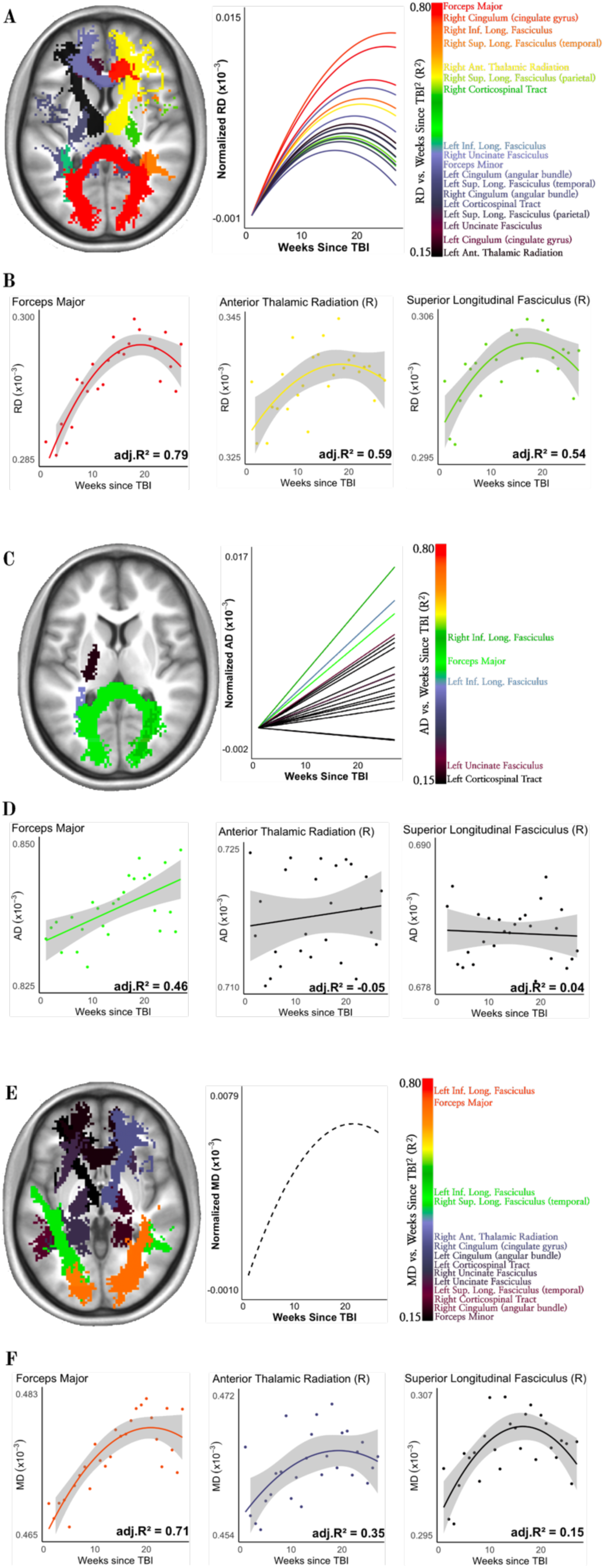
Trajectories of Radial Diffusivity (RD), Axial Diffusivity (AD) and Mean Diffusivity (MD) Changes Over Weeks since TBI. (A) Left: White matter tracts for TBI patient are colored by strength of model (quadratic) fit for RD over time (in weeks) since TBI. Tracts in orange-red indicate a strong relationship (variance explained ∼80%), while tracts in purple-black indicate a weaker relationship (variance explained ∼15%). Right: RD trajectories across all 18 tracts over time, based on the best-fitting quadratic model with random slope effects. RD trajectories are normalized by each tract’s predicted week 1 value to enable simultaneous visualization of all tracts. RD values increase more rapidly in the early weeks post-TBI, before reversing direction, rather than following a simple linear trajectory. (B) Trajectories of RD in three example tracts: forceps major (commissural tract), right anterior thalamic radiation (projection tract), and right superior longitudinal fasciculus (association tract). (C) Left: White matter tracts for TBI patient are colored by model (linear) fit for AD over time (in weeks) since TBI. Right: AD trajectories across all 18 tracts over time, based on the best-fitting linear model with random slope effects. (D) Trajectories of AD in the same three example tracts as in (B). (E) Left: White matter tracts for TBI patient are colored by strength of model (quadratic) fit for MD over time (in weeks) since TBI. Right: MD trajectories across all 18 tracts over time, based on the best-fitting quadratic model with random intercept effects. Given minimal tract-specific variation in slope of MD trajectories, a single broken-line plot illustrates the overall normalized MD trajectory across tracts. (F) Trajectories of MD in the same example tracts as in (B). Across all metrics, tracts outside the 0.15-0.80 range are not represented.

**Figure S2:**
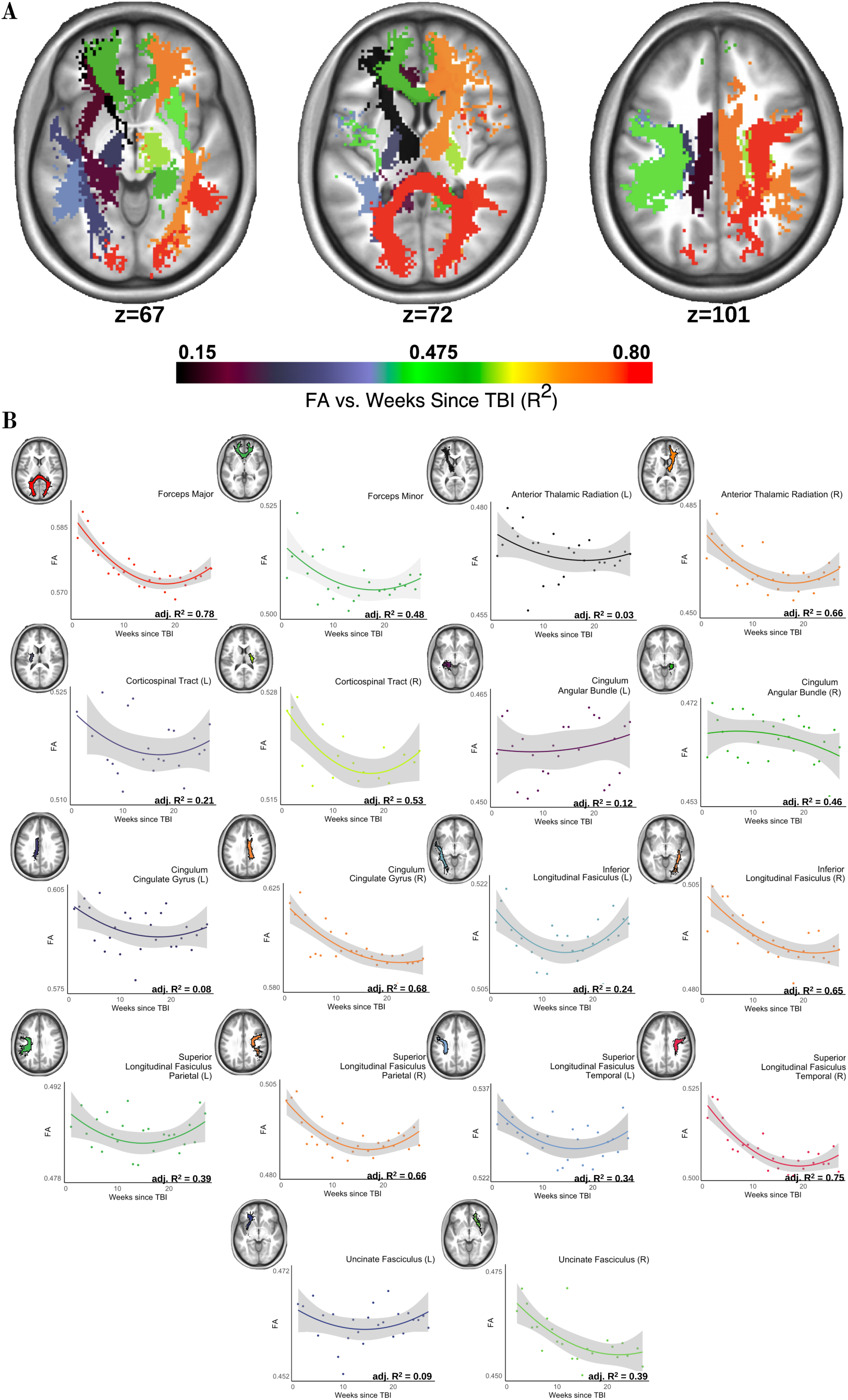
**Fractional Anisotropy (FA) Trajectories Across all Tracts Post-TBI. (**A) Tract- wise R² values (quadratic model) fitting weeks since TBI against FA, visualized in three axial slices. Coordinates indicate Z slice in MNI space. (B) Trajectories of FA in all white matter tracts. In (A) and (B), each white matter tract is color-coded by R² value, representing the proportion of variance explained by the quadratic model.

**Figure S3:**
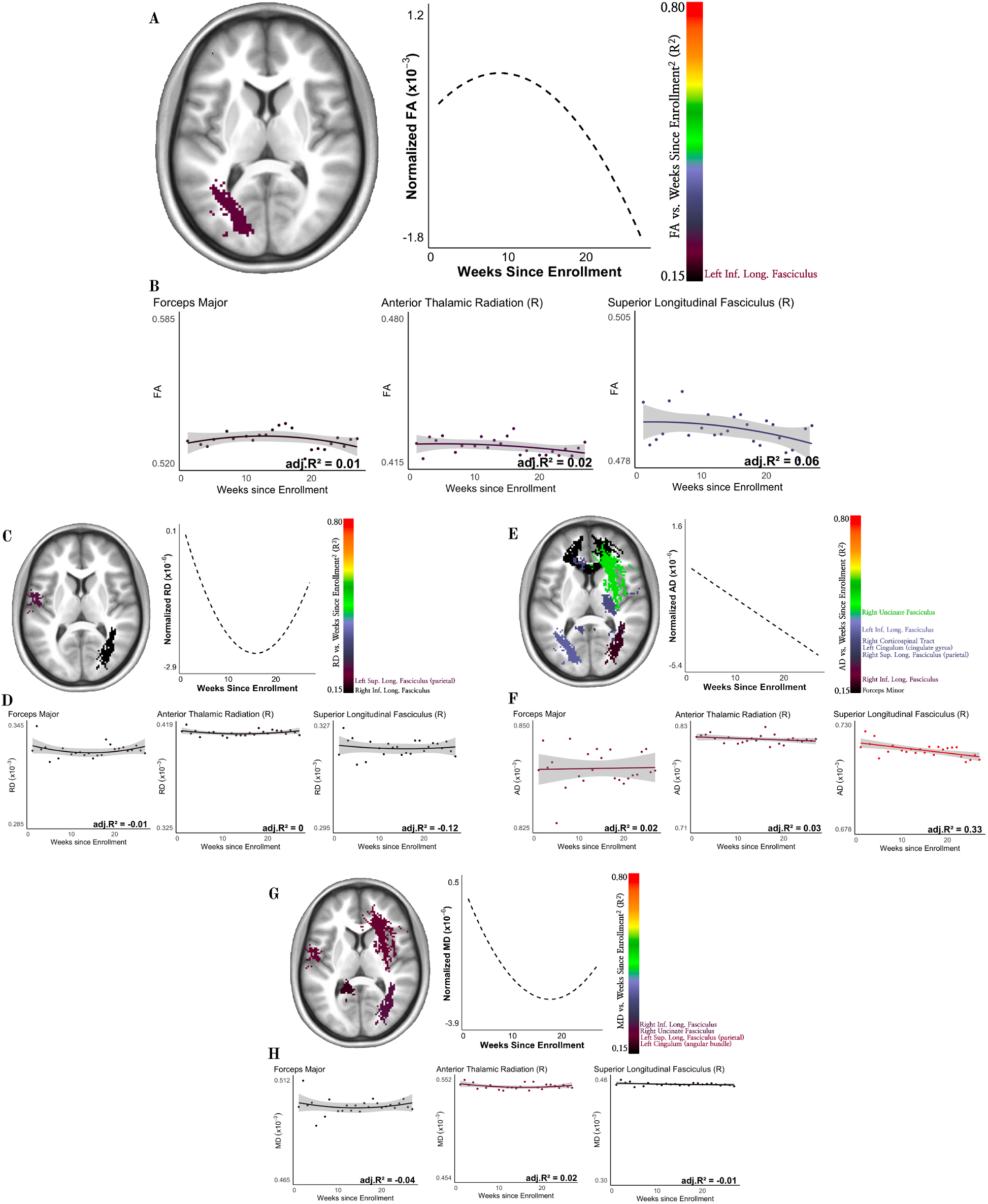
Trajectories of DTI Metrics over Weeks since Enrollment in the Control Participant. (A) White matter tracts for healthy control are colored by strength of model (quadratic) fit for FA over time (in weeks) since enrollment. Right: A single broken-line plot depicts the overall normalized FA trajectory across all 18 tracts over time. (B) Modeled FA trajectories in three representative tracts chosen for direct comparison with the TBI participant (Figure 1): forceps major (commissural tract), right anterior thalamic radiation (projection tract), and right superior longitudinal fasciculus (association tract). (C) Left: White matter tracts for healthy control are colored by strength of model (quadratic) fit for RD over time (in weeks) since enrollment, following the same approach as in (A). Right: a single broken-line plot of overall normalized RD trajectories across all tracts over time. (D) Modeled RD trajectories for the same three example tracts as in (B). (E) Left: White matter tracts for healthy control are colored by strength of model (linear) fit for AD over time (in weeks) since enrollment. Right: a single broken-line plot of overall normalized AD trajectories across all tracts over time. (F) Modeled AD trajectories for the same three example tracts as in (B). (G) Left: White matter tracts for healthy control are colored by strength of model (quadratic) fit for MD over time (in weeks) since enrollment. Right: a single broken-line plot of overall normalized MD trajectories across all tracts over time, (H) Modeled MD trajectories for the same three example tracts as in (B). Unlike the TBI participant, random slopes did not significantly improve model fit for any DTI metric in the control participant, suggesting minimal tract-specific variation in DTI values over time. Instead, DTI metrics in the control participant remained relatively stable, with only minor fluctuations. Across all metrics, tracts outside the 0.15-0.80 range are not represented. Y-axis scales have been adjusted for easy comparison with the TBI patient.

**Figure S4.**
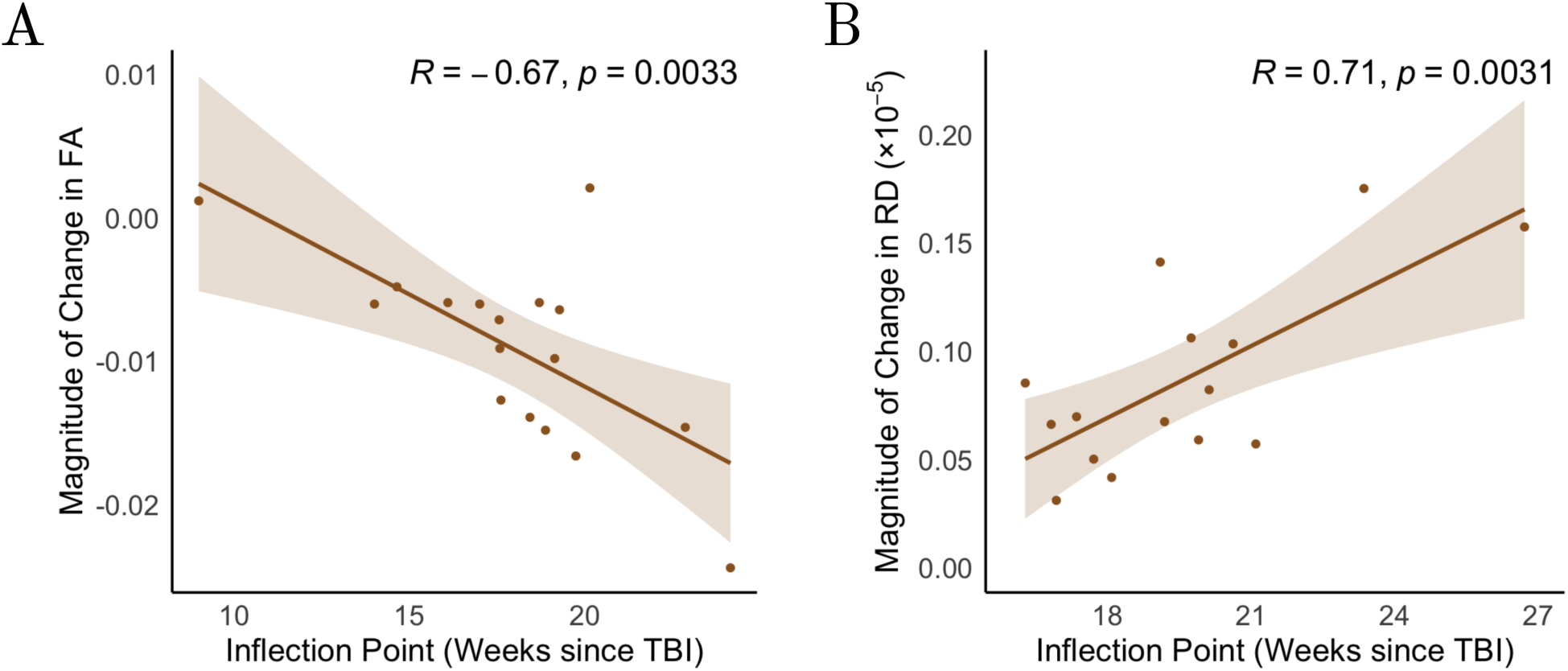
**Relationship Between Inflection Point Timing and Magnitude of Change in FA and RD**. (A) Scatter plot showing the negative correlation between the inflection point timing and the magnitude of change in fractional anisotropy (FA; R = -0.67, p = 0.0033). Later inflection points are associated with greater FA reductions before stabilization. (B) Scatter plot showing the positive correlation between the inflection point timing and the magnitude of change in radial diffusivity (RD; R = 0.71, p = 0.0031). Later inflection points are linked to greater RD increases before stabilization. Shaded areas represent 95% confidence intervals.

**Figure S5.**
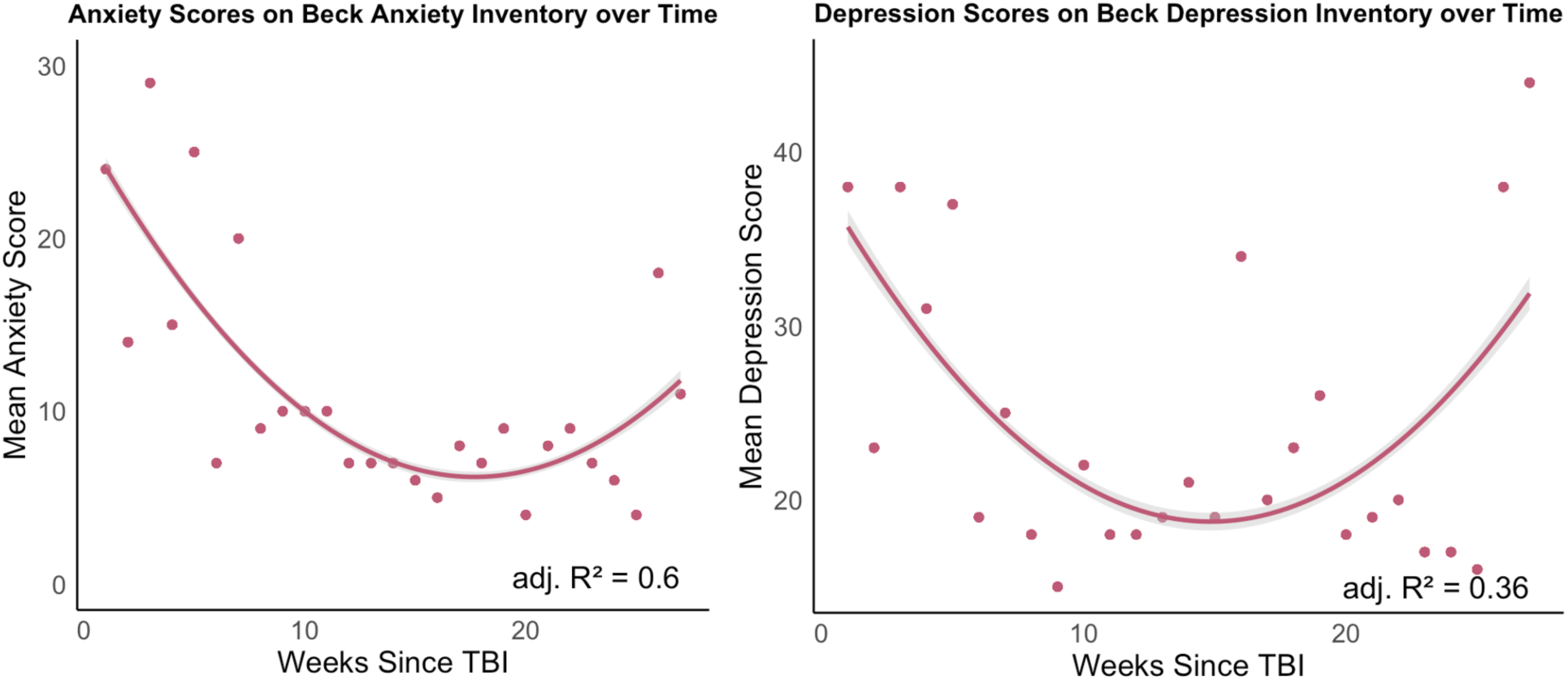
**Trajectories of Anxiety and Depression Symptoms Following TBI**. Mean anxiety (left) and depression (right) scores over 27 weeks post-TBI, assessed using the Beck Anxiety Inventory (Beck, Epstein, Brown, & Steer, R., 1993) and Beck Depression Inventory (Beck, Steer, & Brown, 1996) respectively. Both symptoms followed a quadratic trajectory, with anxiety scores declining sharply before stabilizing, whereas depression scores exhibited a U-shaped pattern, initially improving before later rebounding. Adjusted R² values indicate a stronger fit for anxiety (R² = 0.6) than depression (R² = 0.36), suggesting that anxiety reductions may follow a more predictable course, while depression symptoms may be more variable over time.

**Figure S6.**
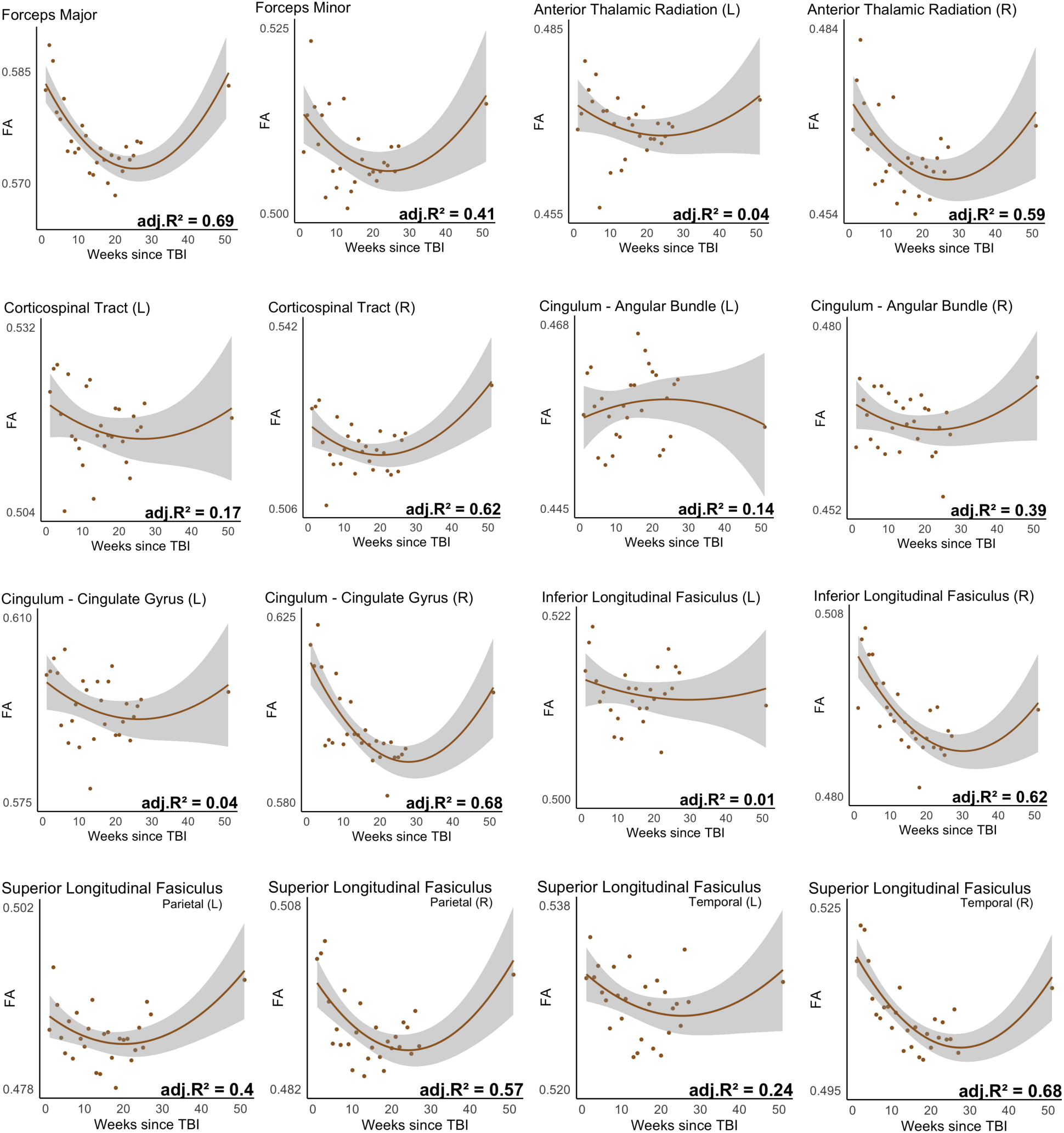
**Fractional Anisotropy (FA) Recovery Trajectories Across White Matter Tracts Post-TBI Up to 1-Year**: Scatterplots illustrate the nonlinear trajectories of FA recovery across major white matter tracts over time (weeks since TBI). Brown lines represent quadratic fits with 95% confidence intervals (gray shading), while adjusted R² values quantify model fit. Right- hemisphere tracts generally exhibit stronger adherence to the quadratic model, as reflected in higher R² values, whereas left-sided tracts show greater variability and weaker model fits, suggesting asymmetric recovery patterns. Notably, the left inferior longitudinal fasciculus (ILF) and left uncinate fasciculus display the largest declines in R² from ∼27 weeks to 1 year, whereas right-sided tracts, including the right ILF and right uncinate, maintain relatively stable R² values, supporting a continued nonlinear trajectory of recovery. These findings reinforce the role of lateralized neurobiological mechanisms in post-TBI white matter repair.

## Notes

### Competing Interest Statement

The authors have declared no competing interest.

## References

1. Abdullah, A. N., Ahmad, A. H., Zakaria, R., Tamam, S., Abd Hamid, A. I., Chai, W. J., … & Abdullah, J. M. (2022). Disruption of white matter integrity and its relationship with cognitive function in non-severe traumatic brain injury. Frontiers in Neurology, 13, 1011304.

2. Amyot, F., Arciniegas, D. B., Brazaitis, M. P., Curley, K. C., Diaz-Arrastia, R., Gandjbakhche, A., … & Stocker, D. (2015). A review of the effectiveness of neuroimaging modalities for the detection of traumatic brain injury. Journal of neurotrauma, 32(22), 1693–1721.

3. Andersson, J. L., & Sotiropoulos, S. N. (2016). An integrated approach to correction for off-resonance effects and subject movement in diffusion MR imaging. Neuroimage, 125, 1063–1078.

4. Armstrong, R. C., Mierzwa, A. J., Marion, C. M., & Sullivan, G. M. (2016). White matter involvement after TBI: Clues to axon and myelin repair capacity. Experimental neurology, 275, 328–333.

5. Barr, D. J., Levy, R., Scheepers, C., & Tily, H. J. (2013). Random effects structure for confirmatory hypothesis testing: Keep it maximal. Journal of memory and language, 68(3), 255–278.

6. Baron, R. M., & Kenny, D. A. (1986). The moderator–mediator variable distinction in social psychological research: Conceptual, strategic, and statistical considerations. Journal of personality and social psychology, 51(6), 1173.

7. Bartels, C., Wegrzyn, M., Wiedl, A., Ackermann, V., & Ehrenreich, H. (2010). Practice effects in healthy adults: a longitudinal study on frequent repetitive cognitive testing. BMC neuroscience, 11, 1–12.

8. Bates, D., Mächler, M., Bolker, B., & Walker, S. (2015). Fitting linear mixed-effects models using lme4. Journal of statistical software, 67, 1–48.

9. Bartoń K (2024). *MuMIn: Multi-Model Inference*. R package version 1.48.4, https://CRAN.R-project.org/package=MuMIn.

10. Beck, A. T., Epstein, N., Brown, G., & Steer, R. (1993). Beck anxiety inventory. Journal of consulting and clinical psychology.

11. Beck, A. T., Steer, R. A., & Brown, G. K. (1996). Beck depression inventory.

12. Bendlin, B. B., Ries, M. L., Lazar, M., Alexander, A. L., Dempsey, R. J., Rowley, H. A., … & Johnson, S. C. (2008). Longitudinal changes in patients with traumatic brain injury assessed with diffusion-tensor and volumetric imaging. Neuroimage, 42(2), 503–514.

13. Braga, R. M., & Buckner, R. L. (2017). Parallel interdigitated distributed networks within the individual estimated by intrinsic functional connectivity. Neuron, 95(2), 457–471.

14. Brickman, A. M., Zimmerman, M. E., Paul, R. H., Grieve, S. M., Tate, D. F., Cohen, R. A., … & Gordon, E. (2006). Regional white matter and neuropsychological functioning across the adult lifespan. Biological psychiatry, 60(5), 444–453.

15. Baum, G. L., Roalf, D. R., Cook, P. A., Ciric, R., Rosen, A. F., Xia, C., … & Satterthwaite, T. D. (2018). The impact of in-scanner head motion on structural connectivity derived from diffusion MRI. Neuroimage, 173, 275–286.

16. Blyth, C. R. (1972). On Simpson’s paradox and the sure-thing principle. Journal of the American Statistical Association, 67(338), 364–366.

17. Bumham, K. P., & Anderson, D. R. (2002). Model selection and multimodel inference: a practical information-theoretic approach. *Spnnger-Veflag, New York*, New York.

18. Carroll, L., Cassidy, J. D., Peloso, P., Borg, J., Von Holst, H., Holm, L., … & Pépin, M. (2004). Prognosis for mild traumatic brain injury: results of the WHO Collaborating Centre Task Force on Mild Traumatic Brain Injury. Journal of rehabilitation medicine, 36(0), 84–105.

19. Catani, M., Dell’Acqua, F., Bizzi, A., Forkel, S. J., Williams, S. C., Simmons, A., … & de Schotten, M. T. (2012). Beyond cortical localization in clinico-anatomical correlation. cortex, 48(10), 1262–1287.

20. Chiang, C. W., Wang, Y., Sun, P., Lin, T. H., Trinkaus, K., Cross, A. H., & Song, S. K. (2014). Quantifying white matter tract diffusion parameters in the presence of increased extra-fiber cellularity and vasogenic edema. Neuroimage, 101, 310–319.

21. Chiou, K. S., Jiang, T., Chiaravalloti, N., Hoptman, M. J., DeLuca, J., & Genova, H. (2019). Longitudinal examination of the relationship between changes in white matter organization and cognitive outcome in chronic TBI. Brain injury, 33(7), 846–853.

22. Cook, J. A., Ramsaya, C. R., & Fayers, P. (2004). Statistical evaluation of learning curve effects in surgical trials. Clinical Trials, 1(5), 421–427.

23. Cragg, J. J., Kramer, J. L., Borisoff, J. F., Patrick, D. M., & Ramer, M. S. (2019). Ecological fallacy as a novel risk factor for poor translation in neuroscience research: A systematic review and simulation study. European journal of clinical investigation, 49(2), e13045.

24. Dosenbach, N. U., Koller, J. M., Earl, E. A., Miranda-Dominguez, O., Klein, R. L., Van, A. N., … & Fair, D. A. (2017). Real-time motion analytics during brain MRI improve data quality and reduce costs. Neuroimage, 161, 80–93.

25. Edlow, B. L., Copen, W. A., Izzy, S., Bakhadirov, K., van der Kouwe, A., Glenn, M. B., … & Wu, O. (2016). Diffusion tensor imaging in acute-to-subacute traumatic brain injury: a longitudinal analysis. BMC neurology, 16, 1–11.

26. Eierud, C., Craddock, R. C., Fletcher, S., Aulakh, M., King-Casas, B., Kuehl, D., & LaConte, S. M. (2014). Neuroimaging after mild traumatic brain injury: review and meta-analysis. NeuroImage: Clinical, 4, 283–294.

27. Figley, C. R., Uddin, M. N., Wong, K., Kornelsen, J., Puig, J., & Figley, T. D. (2022). Potential pitfalls of using fractional anisotropy, axial diffusivity, and radial diffusivity as biomarkers of cerebral white matter microstructure. Frontiers in Neuroscience, 15, 799576.

28. Filley, C. (2012). The behavioral neurology of white matter. Oxford University Press.

29. Fisher, A. J., Medaglia, J. D., & Jeronimus, B. F. (2018). Lack of group-to-individual generalizability is a threat to human subjects research. Proceedings of the National Academy of Sciences, 115(27), E6106–E6115.

30. Forkel, S. J., Friedrich, P., Thiebaut de Schotten, M., & Howells, H. (2022). White matter variability, cognition, and disorders: a systematic review. Brain Structure and Function, 1- 16.

31. Frencham, K. A., Fox, A. M., & Maybery, M. T. (2005). Neuropsychological studies of mild traumatic brain injury: A meta-analytic review of research since 1995. Journal of clinical and experimental neuropsychology, 27(3), 334–351.

32. Galvin, J. E., Tolea, M. I., Moore, C., & Chrisphonte, S. (2020). The Number Symbol Coding Task: A brief measure of executive function to detect dementia and cognitive impairment. PLoS One, 15(11), e0242233.

33. Gennarelli, T. A., Thibault, L. E., Adams, J. H., Graham, D. I., Thompson, C. J., & Marcincin, R. P. (1982). Diffuse axonal injury and traumatic coma in the primate. Annals of Neurology: Official Journal of the American Neurological Association and the Child Neurology Society, 12(6), 564–574.

34. Gratton, C., Kraus, B. T., Greene, D. J., Gordon, E. M., Laumann, T. O., Nelson, S. M., … & Petersen, S. E. (2020). Defining individual-specific functional neuroanatomy for precision psychiatry. Biological psychiatry, 88(1), 28–39.

35. Gratton, C., Nelson, S. M., & Gordon, E. M. (2022). Brain-behavior correlations: Two paths toward reliability. Neuron, 110(9), 1446–1449.

36. Gordon, E. M., Laumann, T. O., Gilmore, A. W., Newbold, D. J., Greene, D. J., Berg, J. J., … & Dosenbach, N. U. (2017). Precision functional mapping of individual human brains. Neuron, 95(4), 791–807.

37. Gordon, E. M., Chauvin, R. J., Van, A. N., Rajesh, A., Nielsen, A., Newbold, D. J., … & Dosenbach, N. U. (2023). A somato-cognitive action network alternates with effector regions in motor cortex. Nature, 617(7960), 351–359.

38. Gyebnár, G., Szabó, Á., Sirály, E., Fodor, Z., Sákovics, A., Salacz, P., … & Csukly, G. (2018). What can DTI tell about early cognitive impairment?–Differentiation between MCI subtypes and healthy controls by diffusion tensor imaging. Psychiatry Research: Neuroimaging, 272, 46–57.

39. Han, D. H., Renshaw, P. F., Dager, S. R., Chung, A., Hwang, J., Daniels, M. A., … & Lyoo, I. K. (2008). Altered cingulate white matter connectivity in panic disorder patients. Journal of psychiatric research, 42(5), 399–407.

40. Harris, N. G., Verley, D. R., Gutman, B. A., & Sutton, R. L. (2016). Bi-directional changes in fractional anisotropy after experiment TBI: Disorganization and reorganization? Neuroimage, 133, 129–143.

41. Holm, S. P., Wolfer, A. M., Pointeau, G. H., Lipsmeier, F., & Lindemann, M. (2022). Practice effects in performance outcome measures in patients living with neurologic disorders–A systematic review. Heliyon, 8(8).

42. Hulkower, M. B., Poliak, D. B., Rosenbaum, S. B., Zimmerman, M. E., & Lipton, M. L. (2013). A decade of DTI in traumatic brain injury: 10 years and 100 articles later. American Journal of Neuroradiology, 34(11), 2064–2074.

43. Jaeger, J. (2018). Digit symbol substitution test: the case for sensitivity over specificity in neuropsychological testing. Journal of clinical psychopharmacology, 38(5), 513–519.

44. Jiang, M., Li, C. L., Zhang, S. Y., Gao, X., & Yang, X. F. (2023). The incidence of brain trauma caused by road injuries: Results from the Global Burden of Disease Study 2019. Injury, 54(12), 110984.

45. Karlsen, R. H., Einarsen, C., Moe, H. K., Håberg, A. K., Vik, A., Skandsen, T., & Eikenes, L. (2019). Diffusion kurtosis imaging in mild traumatic brain injury and postconcussional syndrome. Journal of neuroscience research, 97(5), 568–581.

46. Kievit, R. A., Frankenhuis, W. E., Waldorp, L. J., & Borsboom, D. (2013). Simpson’s paradox in psychological science: a practical guide. Frontiers in psychology, 4, 513.

47. Kim, E., Yoo, R. E., Seong, M. Y., & Oh, B. M. (2022). A systematic review and data synthesis of longitudinal changes in white matter integrity after mild traumatic brain injury assessed by diffusion tensor imaging in adults. European Journal of Radiology, 147, 110117.

48. Kochunov, P., Thompson, P. M., Lancaster, J. L., Bartzokis, G., Smith, S., Coyle, T., … & Fox, P. T. (2007). Relationship between white matter fractional anisotropy and other indices of cerebral health in normal aging: tract-based spatial statistics study of aging. Neuroimage, 35(2), 478–487.

49. Kochunov, P., Williamson, D. E., Lancaster, J., Fox, P., Cornell, J., Blangero, J., & Glahn, D. C. (2012). Fractional anisotropy of water diffusion in cerebral white matter across the lifespan. Neurobiology of aging, 33(1), 9–20.

50. Kraus, M. F., Susmaras, T., Caughlin, B. P., Walker, C. J., Sweeney, J. A., & Little, D. M. (2007). White matter integrity and cognition in chronic traumatic brain injury: a diffusion tensor imaging study. Brain, 130(10), 2508–2519.

51. Kraus, B., Zinbarg, R., Braga, R. M., Nusslock, R., Mittal, V. A., & Gratton, C. (2023). Insights from personalized models of brain and behavior for identifying biomarkers in psychiatry. Neuroscience & Biobehavioral Reviews, 152, 105259.

52. Laumann, T. O., Gordon, E. M., Adeyemo, B., Snyder, A. Z., Joo, S. J., Chen, M. Y., … & Petersen, S. E. (2015). Functional system and areal organization of a highly sampled individual human brain. Neuron, 87(3), 657–670.

53. Laumann, T., Snyder, A., Brown, H., Gott, B., Gordon, E., Barch, D., … & Conway, C. (2023, December). Feasibility of Precision Functional Mapping in Treatment-Resistant Depression. Neuropsychopharmacology, 48, 287–288.

54. Lindsey, H. M., Hodges, C. B., Greer, K. M., Wilde, E. A., & Merkley, T. L. (2023). Diffusion-weighted imaging in mild traumatic brain injury: a systematic review of the literature. Neuropsychology Review, 33(1), 42–121.

55. Ling, J. M., Pena, A., Yeo, R. A., Merideth, F. L., Klimaj, S., Gasparovic, C., & Mayer, A. R. (2012). Biomarkers of increased diffusion anisotropy in semi-acute mild traumatic brain injury: a longitudinal perspective. Brain, 135(4), 1281–1292.

56. Leow, A. D., Zhan, L., Zhu, S., Hageman, N., Chiang, M. C., Barysheva, M., … & Thompson, P. M. (2009, June). White matter integrity measured by fractional anisotropy correlates poorly with actual individual fiber anisotropy. In 2009 *IEEE International Symposium on Biomedical Imaging: From Nano to Macro* (pp. 622-625). IEEE.

57. Lynch, C. J., Elbau, I. G., Ng, T., Ayaz, A., Zhu, S., Wolk, D., … & Liston, C. (2024). Frontostriatal salience network expansion in individuals in depression. Nature, 633(8030), 624–633.

58. Maas, A. I., Menon, D. K., Manley, G. T., Abrams, M., Åkerlund, C., Andelic, N., … & Zemek, R. (2022). Traumatic brain injury: progress and challenges in prevention, clinical care, and research. The Lancet Neurology, 21(11), 1004–1060.

59. Mattoni, M., Fisher, A. J., Gates, K. M., Chein, J., & Olino, T. M. (2025). Group-to-Individual Generalizability and Individual-Level Inferences in Cognitive Neuroscience. Neuroscience & Biobehavioral Reviews, 106024.

60. Mayer, A. R., Ling, J., Mannell, M. V., Gasparovic, C., Phillips, J. P., Doezema, D., … & Yeo, R. (2010). A prospective diffusion tensor imaging study in mild traumatic brain injury. Neurology, 74(8), 643–650.

61. Mesaros, S., Rocca, M. A., Kacar, K., Kostic, J., Copetti, M., Stosic-Opincal, T., … & Filippi, M. (2012). Diffusion tensor MRI tractography and cognitive impairment in multiple sclerosis. Neurology, 78(13), 969–975.

62. Michon, K. J., Khammash, D., Simmonite, M., Hamlin, A. M., & Polk, T. A. (2022). Person-specific and precision neuroimaging: Current methods and future directions. NeuroImage, 263, 119589.

64. Murphy, M. L., & Frodl, T. (2011). Meta-analysis of diffusion tensor imaging studies shows altered fractional anisotropy occurring in distinct brain areas in association with depression. Biology of mood & anxiety disorders, 1, 1–12.

65. Næss-Schmidt, E. T., Blicher, J. U., Eskildsen, S. F., Tietze, A., Hansen, B., Stubbs, P. W., … & Nielsen, J. F. (2017). Microstructural changes in the thalamus after mild traumatic brain injury: a longitudinal diffusion and mean kurtosis tensor MRI study. Brain injury, 31(2), 230–236.

66. Nakayama, N., Okumura, A., Shinoda, J., Yasokawa, Y. T., Miwa, K., Yoshimura, S. I., & Iwama, T. (2006). Evidence for white matter disruption in traumatic brain injury without macroscopic lesions. *Journal of Neurology*, Neurosurgery & Psychiatry, 77(7), 850–855.

67. Narayana, P. A., Yu, X., Hasan, K. M., Wilde, E. A., Levin, H. S., Hunter, J. V., … & McCarthy, J. J. (2015). Multi-modal MRI of mild traumatic brain injury. NeuroImage: Clinical, 7, 87–97.

68. Naselaris, T., Allen, E., & Kay, K. (2021). Extensive sampling for complete models of individual brains. Current Opinion in Behavioral Sciences, 40, 45–51.

69. Newbold, D. J., Laumann, T. O., Hoyt, C. R., Hampton, J. M., Montez, D. F., Raut, R. V., … & Dosenbach, N. U. (2020). Plasticity and spontaneous activity pulses in disused human brain circuits. Neuron, 107(3), 580–589.

70. Niogi, S. N., & Mukherjee, P. (2010). Diffusion tensor imaging of mild traumatic brain injury. The Journal of head trauma rehabilitation, 25(4), 241–255.

71. Palacios, E. M., Owen, J. P., Yuh, E. L., Wang, M. B., Vassar, M. J., Ferguson, A. R., … & Track-TBI Investigators. (2020). The evolution of white matter microstructural changes after mild traumatic brain injury: a longitudinal DTI and NODDI study. Science advances, 6(32), eaaz6892.

72. Pham, L., Harris, T., Varosanec, M., Morgan, V., Kosa, P., & Bielekova, B. (2021). Smartphone-based symbol-digit modalities test reliably captures brain damage in multiple sclerosis. NPJ digital medicine, 4(1), 36.

73. Pierpaoli, C., Barnett, A., Pajevic, S., Chen, R., Penix, L., Virta, A., & Basser, P. (2001). Water diffusion changes in Wallerian degeneration and their dependence on white matter architecture. Neuroimage, 13(6), 1174–1185.

74. Preacher, K. J., & Hayes, A. F. (2008). Asymptotic and resampling strategies for assessing and comparing indirect effects in multiple mediator models. Behavior research methods, 40(3), 879–891.

75. Preziosa, P., Pagani, E., Meani, A., Marchesi, O., Conti, L., Falini, A., … & Filippi, M. (2023). NODDI, diffusion tensor microstructural abnormalities and atrophy of brain white matter and gray matter contribute to cognitive impairment in multiple sclerosis. Journal of Neurology, 270(2), 810-823.

76. Pritschet, L., Santander, T., Taylor, C. M., Layher, E., Yu, S., Miller, M. B., … & Jacobs, E. G. (2020). Functional reorganization of brain networks across the human menstrual cycle. Neuroimage, 220, 117091.

77. Pritschet, L., Taylor, C. M., Cossio, D., Faskowitz, J., Santander, T., Handwerker, D. A., … & Jacobs, E. G. (2024). Neuroanatomical changes observed over the course of a human pregnancy. Nature Neuroscience, 27(11), 2253–2260.

78. Roberts, R. E., Anderson, E. J., & Husain, M. (2013). White matter microstructure and cognitive function. The Neuroscientist, 19(1), 8–15.

79. Robinson, W. S. (2009). Ecological correlations and the behavior of individuals. International journal of epidemiology, 38(2), 337–341.

80. Rosenbaum, S. B., & Lipton, M. L. (2012). Embracing chaos: the scope and importance of clinical and pathological heterogeneity in mTBI. Brain imaging and behavior, 6, 255–282.

81. Schmahmann, J. D., Smith, E. E., Eichler, F. S., & Filley, C. M. (2008). Cerebral white matter: neuroanatomy, clinical neurology, and neurobehavioral correlates. Annals of the New York Academy of Sciences, 1142(1), 266–309.

82. Schretlen, D. J., & Shapiro, A. M. (2003). A quantitative review of the effects of traumatic brain injury on cognitive functioning. International review of psychiatry, 15(4), 341–349.

83. Schwartz, S. (1994). The fallacy of the ecological fallacy: the potential misuse of a concept and the consequences. American journal of public health, 84(5), 819–824.

84. Sexton, C. E., Mackay, C. E., & Ebmeier, K. P. (2009). A systematic review of diffusion tensor imaging studies in affective disorders. Biological psychiatry, 66(9), 814–823.

85. Shenton, M. E., Hamoda, H. M., Schneiderman, J. S., Bouix, S., Pasternak, O., Rathi, Y., … & Zafonte, R. (2012). A review of magnetic resonance imaging and diffusion tensor imaging findings in mild traumatic brain injury. Brain imaging and behavior, 6, 137–192.

86. Siegel, J. S., Subramanian, S., Perry, D., Kay, B. P., Gordon, E. M., Laumann, T. O., … & Dosenbach, N. U. (2024). Psilocybin desynchronizes the human brain. Nature, 632(8023), 131–138.

87. Smith, A. (1973). Symbol digit modalities test. The clinical neuropsychologist.

88. Smith, S. M., Jenkinson, M., Woolrich, M. W., Beckmann, C. F., Behrens, T. E., Johansen-Berg, H., … & Matthews, P. M. (2004). Advances in functional and structural MR image analysis and implementation as FSL. Neuroimage, 23, S208–S219.

89. Teasdale, G., & Jennett, B. (1974). Assessment of coma and impaired consciousness. A Practical Scale Lancet 2: 81*–*84.

90. Tingley, D., Yamamoto, T., Hirose, K., Keele, L., & Imai, K. (2014). Mediation: R package for causal mediation analysis. Journal of statistical software, 59, 1–38.

91. Wagner, C. H. (1982). Simpson’s paradox in real life. The American Statistician, 36(1), 46–48.

92. Wallace, E. J., Mathias, J. L., & Ward, L. (2018). Diffusion tensor imaging changes following mild, moderate and severe adult traumatic brain injury: a meta-analysis. Brain imaging and behavior, 12, 1607–1621.

93. Wang, Y., Wang, Q., Haldar, J. P., Yeh, F. C., Xie, M., Sun, P., … & Song, S. K. (2011). Quantification of increased cellularity during inflammatory demyelination. Brain, 134(12), 3590–3601.

94. Wang, X., Cusick, M. F., Wang, Y., Sun, P., Libbey, J. E., Trinkaus, K., … & Song, S. K. (2014). Diffusion basis spectrum imaging detects and distinguishes coexisting subclinical inflammation, demyelination and axonal injury in experimental autoimmune encephalomyelitis mice. NMR in Biomedicine, 27(7), 843–852.

95. Wang, Y., Sun, P., Wang, Q., Trinkaus, K., Schmidt, R. E., Naismith, R. T., … & Song, S. K. (2015). Differentiation and quantification of inflammation, demyelination and axon injury or loss in multiple sclerosis. Brain, 138(5), 1223–1238.

96. Wang, S., Peterson, D. J., Wang, Y., Wang, Q., Grabowski, T. J., Li, W., & Madhyastha, T. M. (2017). Empirical comparison of diffusion kurtosis imaging and diffusion basis spectrum imaging using the same acquisition in healthy young adults. Frontiers in neurology, 8, 118.

97. Wang, Q., Wang, Y., Liu, J., Sutphen, C. L., Cruchaga, C., Blazey, T., … & Benzinger, T. L. (2019a). Quantification of white matter cellularity and damage in preclinical and early symptomatic Alzheimer’s disease. NeuroImage: Clinical, 22, 101767.

98. Wang, Q., Pérez-Carrillo, G. J. G., Ponisio, M. R., LaMontagne, P., Dahiya, S., Marcus, D. S., … & Wang, Y. (2019b). Heterogeneity Diffusion Imaging of gliomas: Initial experience and validation. PloS one, 14(11), e0225093.

99. Wilde, E. A., Li, X., Hunter, J. V., Narayana, P. A., Hasan, K., Biekman, B., … & Levin, H. S. (2016). Loss of consciousness is related to white matter injury in mild traumatic brain injury. Journal of neurotrauma, 33(22), 2000–2010.

100. Veeramuthu, V., Hariri, F., Narayanan, V., Tan, L. K., Ramli, N., & Ganesan, D. (2016). Microstructural change and cognitive alteration in maxillofacial trauma and mild traumatic brain injury: A diffusion tensor imaging study. Journal of Oral and Maxillofacial Surgery, 74(6), 1197–e1.

101. Yendiki, A., Panneck, P., Srinivasan, P., Stevens, A., Zöllei, L., Augustinack, J., … & Fischl, B. (2011). Automated probabilistic reconstruction of white-matter pathways in health and disease using an atlas of the underlying anatomy. Frontiers in neuroinformatics, 5, 23.

102. Zhang, Y., Brady, M., & Smith, S. (2001). Segmentation of brain MR images through a hidden Markov random field model and the expectation-maximization algorithm. IEEE transactions on medical imaging, 20(1), 45-57.

103. Zhang, L., Zhang, Y., Li, L., Li, Z., Li, W., Ma, N., … & Lu, G. (2011). Different white matter abnormalities between the first-episode, treatment-naive patients with posttraumatic stress disorder and generalized anxiety disorder without comorbid conditions. Journal of affective disorders, 133(1-2), 294–299.

